# Sequence-Specific Installation of Aryl Groups in RNA via DNA-Catalyst Conjugates

**DOI:** 10.1101/2025.09.04.674272

**Authors:** Sumon Pratihar, Wenrui Zhong, Sheng Feng, Sayantan Chatterjee, Eric T. Kool

## Abstract

Installing functional groups at specific sites in existing RNA molecules remains a challenge for modification, labeling, and therapeutic strategies. Here we describe the use of DNA oligonucleotides carrying a catalytic amine group to effect the aqueous S_N_Ar arylation of 2′-OH groups at sequence-complementary sites in RNAs. Chloro-pyrimidine electrophiles are shown to react with amino-DNA conjugates, resulting in a proposed transient ammonium aryl intermediate that can react with RNA near the DNA binding site, delivering the heterocycle to the RNA in high yields. In a test of utility, we construct an aryl electrophile carrying an azide group, and apply this strategy to fluorescently label messenger RNAs locally at the polyA tail. We also employ the approach to direct *in vitro* arylation in the coding region of a messenger RNA, knocking down protein expression selectively in the presence of another coding RNA. This sequence-directed catalytic strategy enables multiple applications in RNA labeling and modification.

## Introduction

RNA modification with probes is important for understanding the basic structural dynamics, localization, expression, and functions of the biopolymer. Chemical modification can also be enabling for altering the biological properties of therapeutic RNAs, including delivery, immunogenicity, and intracellular lifetime.^1^ General labelling of RNAs throughout the polymer, such as with fluorescent dyes, is readily available via incorporation of labelled nucleotides during transcription.^2^ However, the presence of labels at many positions in longer RNAs loses the precision of localized interrogation of structure and interaction, and can block biological function where locally unmodified structure is required. Site-specific RNA modification can overcome this issue, and has been widely applied to short RNAs that are chemically synthesized. Ligation-based methods have been reported for locally modifying RNAs that are too long for chemical synthesis,^3^ however they can be laborious and remain to be widely adopted.

A potential answer to these challenges is the development of methods for post-synthetic RNA modification.^4-5^ Such approaches, if localized to specific sites in existing RNAs, can simplify the targeted modification of RNA post-transcriptionally while avoiding the challenging synthesis of nucleotides. Efficient site-specific modification strategies have the potential to tune the cellular lifetime of RNA, alter its local conformation, unravel its utility as drug target, and enable potential implications in gene knockdown.

Biochemical approaches for postsynthetic RNA modification have made use of engineered protein or nucleic acid enzymes. For example, the use of a bacterial tRNA-modifying enzyme with specific modified substrates enables the postsynthetic delivery of labels to guanosine nucleobases in tRNA-mimicking loops engineered into RNA.^6^ The method requires prior RNA engineering to contain reactive sites, and thus is not applicable to native RNAs. More recently, DNAzyme-based strategies targeting the 2’-OH group of RNA have demonstrated considerable success in local modification of non-engineered RNAs; using the 2’-OH group as the site of modification offers the possibility of widespread application, since the group appears at every position of RNA. Employing this strategy, Silverman and Höbartner have described RNA-modifying DNAzymes consisting of a catalytic DNA sequence containing a binding pocket for a nucleotide reagent.^7-9^ Although elegant, such enzymatic approaches exhibit sequence restrictions and preferences that limit generality significantly, and employ modified nucleotides with nontrivial syntheses.

In a different approach, DNA complementarity can be employed to direct RNA modification locally defined by the sequence in the induced-loop methods RAIL and TRAIL.^10,11^ In this strategy, the nontargeted nucleotides in an RNA being targeted are masked by hybridization with complementary DNA, leaving only the desired unmasked nucleotide accessible for reaction at 2’-OH with acylimidazole-based reagents. The strategy is site-specific, made possible by blocking all other ribonucleotides in a strand. However, the method is limited by the large number of DNA oligonucleotides required for blocking very long RNAs (*e*.*g*. > 2000 nt), and is not practical for modifying a specific RNA in a mixture of transcripts.

A potential solution to these challenges lies in the use of small DNA conjugates to deliver groups locally to RNA. In early approaches, DNA oligonucleotides with 6-thioguanine, wherein a reactive moiety is covalently appended to the sulfur atom, were explored for Michael addition of an α,β-unsaturated diketone group selectively to the N4 of proximal cytosines in the complementary RNA.^12^ 6-Thioguanine in DNA was further adapted for site-specific acetylation of the 2′-OH groups in RNA at pH 10, preferentially reacting at cytosine and uridine residues.^13^ More recently, nucleoside derivatives appended with imidazole, pyridine, and pyrimidines were tested in site-specific acetylation of 2’-OH in a 16-mer RNA using acetic anhydride as the electrophile.^14^ While significant advances, none of these approaches were tested with long RNAs nor in mixtures of RNAs where off-target nonspecific reactivity of highly electrophilic reagents, or RNA instability due to basic conditions, may interfere.

Here, we develop a strategy to address these challenges, aiming to execute strategic RNA modification of long biologically active RNAs at desired locations in an unbiased manner under mild conditions. This approach utilizes a DNA oligonucleotide carrying a nucleophilic amine to promote site-selective S_N_Ar reaction at RNA 2′-OH groups proximal to the catalytic group (Figure 1A). The use of aryl electrophiles with low inherent reactivity both to RNA and to water facilitates selective reaction in long RNAs, and enables function with multiple RNAs present. Given the ready modifiability of aryl reactants, we anticipate that the strategy can be applied to a broad range of label types, and may also suggest future strategies for specific gene knockdown.

**Figure 1.**
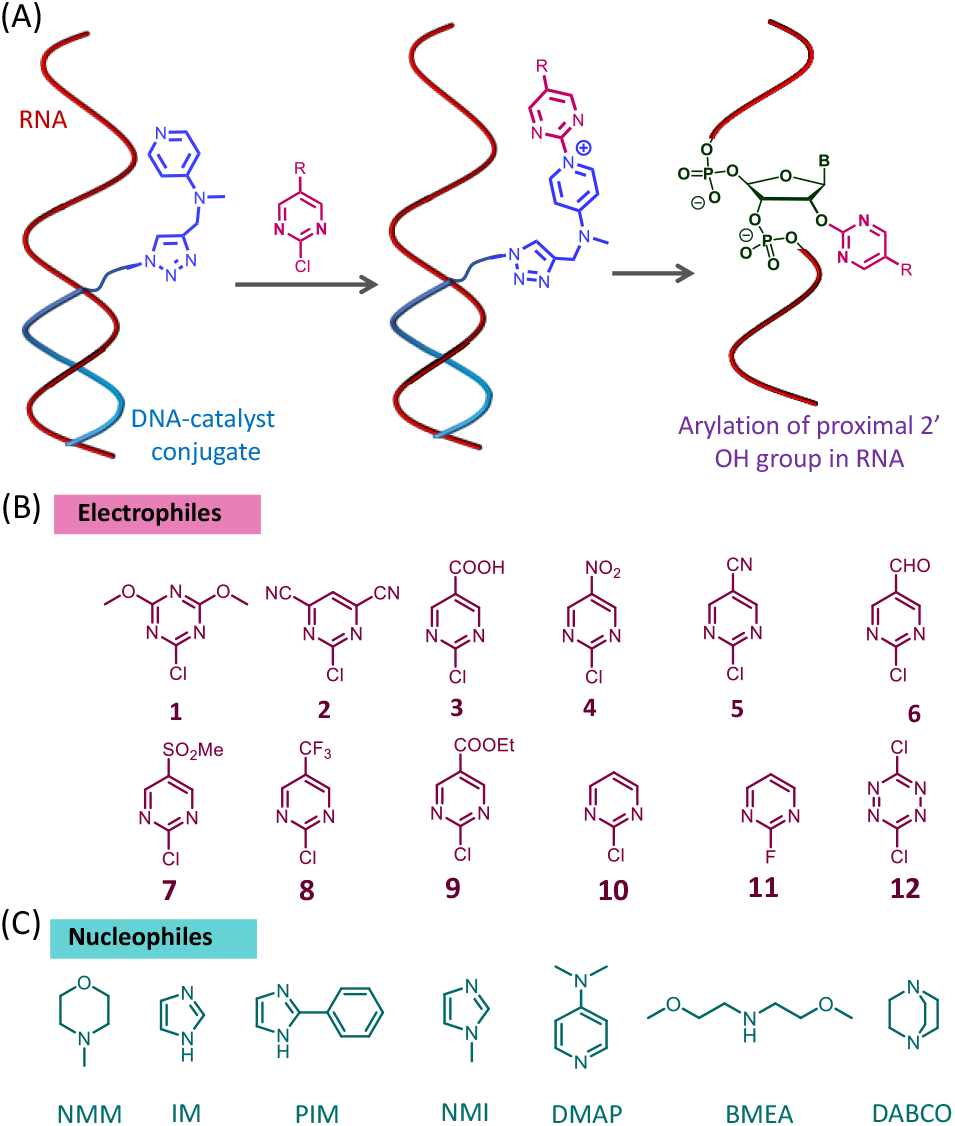
Mechanisms and structures for DNA-directed local RNA arylation at 2’-OH groups. (A) Scheme for employing a DNA-amine conjugate to activate and locally deliver a 2’OH-reactive electrophile to a complementary site in RNA. The DNA oligonucleotide carries a nucleophilic amine that can act as a catalyst, forming a transient activated adduct with a reagent that otherwise reacts poorly with RNA. The activated adduct is delivered at a high local concentration to the RNA nucleotides in proximity of the DNA end. (B) Range of aryl electrophiles tested; (C) Candidate amine nucleophilic catalysts tested.

## Results and Discussion

Recent studies have demonstrated the unusually high nucleophilic reactivity of the 2’-OH group of RNA for high-yield modification, and have characterized multiple classes of electrophiles that can react there efficiently.^15^ While acylation is the most common mechanism, it was found recently that S_N_Ar reactions also can be carried out at the 2’-OH group of RNA, employing aromatic heterocycles bearing quaternary ammonium leaving groups such as N-methylmorpholinium.^16^ That work also described the observation that aryl chlorides showed much lower RNA reactivity than did the corresponding ammonium species. Here, we envisioned the possibility that an RNA-reactive quaternary ammonium species might be formed transiently in aqueous solution by reaction of aryl halide with a nucleophilic amine *in situ* (Figure 1). If the nucleophilic amine were conjugated to a DNA oligonucleotide, this might enable sequence-specific delivery of the arylammonium group to RNA, programmed by the sequence of the DNA.

To be successful, this strategy requires the identification of a nucleophilic tertiary amine that can form a transient adduct with an aryl halide in water, and that this ammonium adduct is highly RNA-reactive. At the same time, the aryl halide, which would be used in excess, must show low RNA reactivity on its own to avoid unwanted off-target effects. To approach this, we examined a series of aromatic heterocyclic electrophiles (Figure 1B) in combination with tertiary amines (Figure 1C) to find the most-efficient electrophile/leaving group pair for selective 2’-OH modification. Examples of S_N_Ar reactions promoted by nucleophilic catalysis are rare in the literature;^17,18^ for nonoenzymatic cases we are aware of a single prior report of a nucleophile-promoted reaction which was carried out in organic solvent at elevated temperature.^17^

To start, we explored the nucleophilic amines (Figure 1C), evaluating their efficiency in forming the corresponding quaternary ammonium adducts with a chlorotriazine electrophile. N-methylmorpholine (NMM), Imidazole (IM), 2-phenylimidazole (PIM), N-methylimidazole (NMI), dimethylaminopyridine (DMAP), bis(methoxyethyl)amine (BMEA) and diazabicyclooctane (DABCO) were reacted with the triazine chloride **1** in tetrahydrofuran as described previously for trimethylamine.^17^ DMAP and NMM gave quantitative yields of the corresponding arylammonium adducts, while NMI also gave excellent yields after extended time (see SI for NMR spectra of selected adducts). Given that our intended application is for aqueous solution, we performed time course measurements of reaction of **1** with the three most reactive nucleophiles in water, and the results confirmed 65-75% reaction in two hours for DMAP and NMM (Figure 2A), with slower reaction by NMI. We anticipated that successful generation of the ammonium arylating reagent *in situ* could engender significant leverage both by rendering them potentially catalytic, and by enabling sequence-directed delivery.

**Figure 2.**
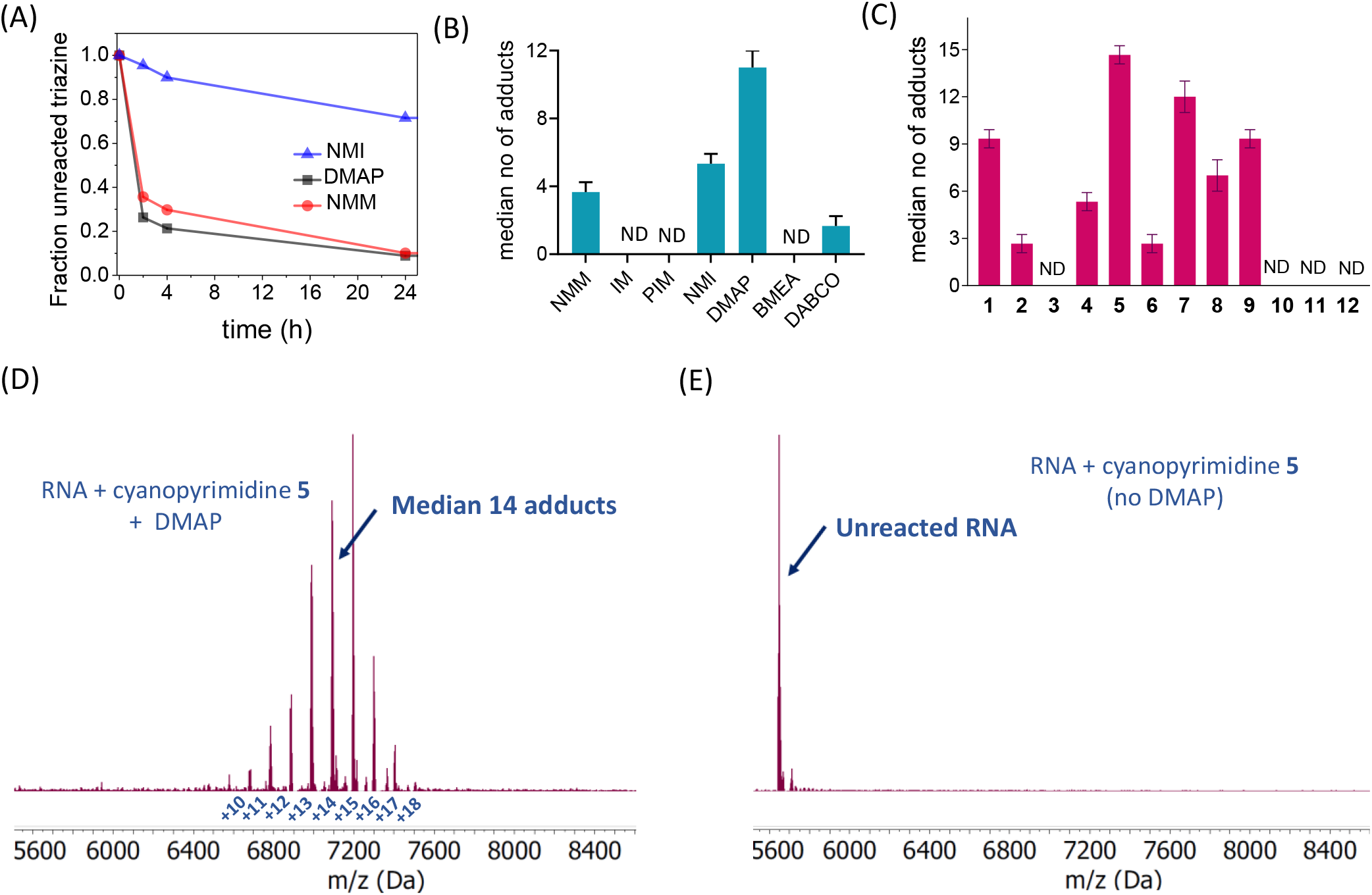
Amine nucleophiles promote the reaction of aryl halides with RNA. (A) Time courses of aqueous reaction of NMI, NMM, and DMAP with triazine chloride **1**; (B) Plot of RNA yields (median number of adducts in a short test RNA) with varied amines (reacting with triazine chloride **1** *in situ*); (C) Comparison of aryl halides **1**-**12** (Figure 1B), shown by graph of RNA adduct yields with the different aryl species in the presence of DMAP (ND = no adducts detected); (D) Representative mass spectra of 18 nt single-stranded RNA (MW 5642 Da) reacted with cyanopyrimidine 5 (100 mM) in the presence of DMAP (100 mM) and (E) without the catalyst. See text for conditions. For (B,C), data show averages from 3 replicates; error bars show standard deviations.

Next, we evaluated the reactivity of the *in situ*-generated ammonium-aryl adducts with a short single-stranded RNA sequence (18 nt) in pH 7.5 MOPS buffer, with 6 mM MgCl_2_, 100 mM NaCl at 37 °C for 24 h. MALDI-TOF mass spectrometric analysis of the reacted RNA revealed the highest reactivity with DMAP (an average of 11 adducts observed), followed by NMI (5.3 adducts) and NMM (3.6 adducts) (Figures 2B, S1) and the overall rank order DMAP > NMM > NMI > DABCO >> BMEA ∼ PIM). No discernible adduct formation was seen with the last two nucleophiles. This reactivity with RNA correlated with the trend of the reactivity of corresponding amine toward the aryl chloride (Figures 2A,B). Given the facile reactivity of DMAP with the aryl chloride and the high reactivity of the DMAP-aryl adduct with RNA, this emerged as a nucleophile of choice, together with NMI and NMM as potential alternatives. Also noteworthy was the fact that the previously unknown DMAP-aryl adduct displayed yet higher RNA reactivity than the recently described NMM adduct (Figures 2B, S2).^16^

Next, we explored electrophiles, testing a range of electron-deficient aromatic halides to arylate an 18 nt RNA in the presence and absence of DMAP, evaluating the RNA reactivity of each DMAP-electrophile combination (Figure 2C, Figure S3). In principle, an ideal reagent would show maximal enhancement by DMAP, while having minimal RNA reactivity on its own. For aqueous solubility we focused on pyrimidine, triazine or tetrazine aryl electrophiles bearing chloride or fluoride leaving groups. For these experiments, we again relied on *in situ* formation of ammonium-aryl adducts, mixing aryl halides 1:1 with DMAP (100 mM each) and RNA (10 μM). This would serve as a functional test not only of the reactivity of the electrophiles, but also of the corresponding ammonium species’ reactivity with RNA; this *in situ* formation is required for our DNA-directed strategy to succeed.

The results revealed that while good conversion of RNA was observed with multiple aryl chlorides, 5-cyanopyrimidine yielded the greatest number of RNA adducts (Figures 2D and S3). Control experiments with DNA (which lacks 2’-OH groups) confirmed 2’-OH as the site of reaction on RNA (Figure S4) as seen previously.^16^ Notably, in the absence of DMAP, minimal reactions were observed with the RNA even at 100 mM pyrimidine chloride, indicating high responsiveness to DMAP but low background reactivity (Figures 2E, S5). The results demonstrate the critical role of DMAP as an effective nucleophilic promoter for the arylation reaction with RNA. Although DMAP has been employed recently to enhance reactivity of imidazole carbamate reagents with RNA,^19, 15^ these new results demonstrated the previously unknown RNA reactivity of aryl groups using DMAP adducts formed *in situ*. We expect that this DMAP-assisted reactivity may also be applied beyond RNA, to protein labelling schemes that involve S_N_Ar chemistry.^20^

Encouraged by the excellent RNA reaction efficiency of the *in situ*-generated arylating agents, we proceeded to test the DNA-amine-based strategy to deliver the arylating groups at targeted locations in RNA. We designed short complementary DNA oligonucleotides (15 nt) bearing a commercial azide linker at its 5′ end connected via amide linkage (Figures 3A, S6). One-step Cu(I)-mediated reaction of the azide modified DNA with alkyne-functionalized amine nucleophiles resulted in DNA conjugates bearing the amines of interest. The DNA conjugation reactions proceeded in 90-97% yields as analyzed by mass spectrometry (Figure S7).

**Figure 3.**
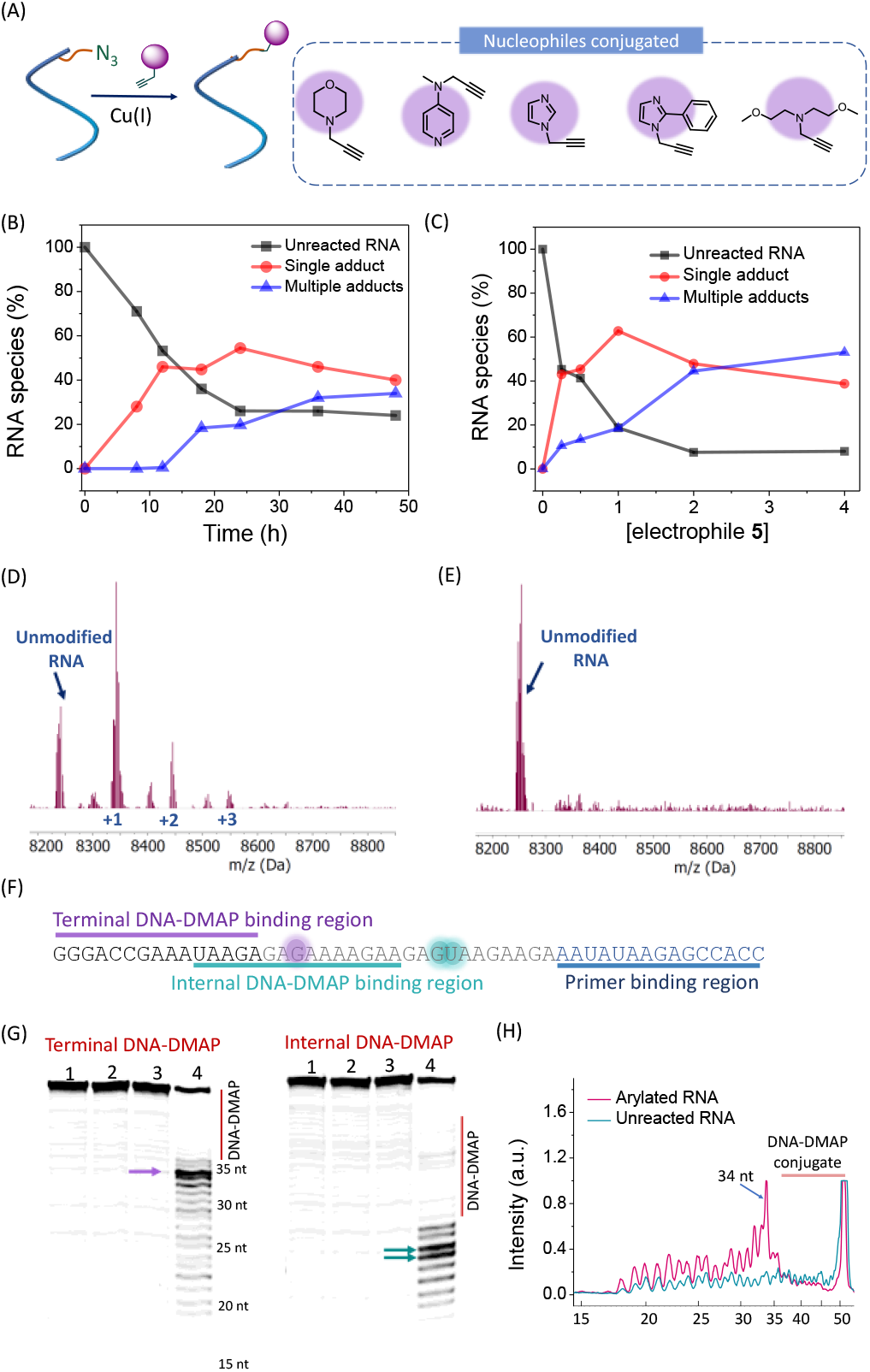
Tests of DNA-directed RNA arylation. (A) Chemical structures of the DNA-amine conjugates tested. (B) Time course of yields of arylation adducts of RNA in presence of 2 mM cyanopyrimidine **5** over 48 h; (C) RNA yields under varied concentrations of **5** (250 µM, 500 µM, 1 mM, 2 mM, 4 mM); (D) and (E) Representative mass spectra of 25 nt RNA reaction products in the presence of a complementary DNA +/-conjugated amine); (F) Representation of the 51 nt RNA sequence used for primer extension and the positions of selective arylation (highlighted in purple/cyan); (G) PAGE gel analysis after cDNA primer extension showing localized modification within the 51nt RNA near the position of the amine conjugate at two different positions in the RNA. Lane 1: RNA, 2: RNA + Cyanopyrimidine **5**, 3: RNA + DNA-DMAP, 4: RNA + DNA-DMAP +Cyanopyrimidine **5**. (H) Quantification band intensities of the lanes 3 and 4 of the gel for reaction with terminal DNA-DMAP conjugate (left); the cDNA lengths (in nucleotides) are given on the x-axis. Uncropped gel image is provided in the Supporting Information.

To attempt sequence-directed RNA modification reactions, we hybridized the 15mer amine-conjugated DNAs (Figure 3A) with a 25 nt RNA, and incubated the complexes with the cyanopyrimidine-based electrophile **5** at varied concentrations in MOPS buffer (pH 7.5) at 37 ^°^C. After 24 h, the imidazole-DNA conjugate provided minimal RNA modification, while the morpholine analog showed moderate reactivity (23 % of a single arylation product, Figure S8). In contrast, the DMAP-DNA conjugate provided good conversion to mono-(47%) and di-arylated (8%) RNA. Employing mass spectrometric analysis, we optimized reaction time (Figure 3B) and aryl chloride concentration (Figure 3C) to achieve highest conversion to singly arylated RNA. We found that 2 mM of the cyanopyrimidine **5** in pH 7.5 buffer yielded 78 % conversion of the RNA, with monoarylation predominant (69% mono-, 9% di-; Figure 3D). Controls employing the same DNA oligonucleotide lacking the DMAP displayed little or no reactivity under identical reaction conditions (Figure 3E). The *in situ* formation of the proposed DNA-ammonium active intermediate proposed to be responsible for driving this reaction was separately confirmed by MALDI-TOF analysis (Figure S9). Effective DNA-directed arylation of RNA was also achieved with electrophile **7** (Figure S10).

Under these conditions, only 10 μM of the DMAP group is present for effective arylation, compared to 2-5 mM necessary for the non-DNA-directed reaction with 18 nt RNA (Figure S11). This demonstrates the high reactivity conferred by the DMAP-DNA conjugate, increasing the effective local concentration of the reactive species by *ca*. 500-fold. The formation of the multiarylation products at longer times suggests the potentially catalytic role of DMAP, wherein the DNA-linked DMAP sequentially reacts with the aryl donor and efficiently delivers it to RNA, regenerating the free DMAP group for further rounds of reaction.

To assess the site-selectivity of the RNA arylation promoted by a DNA-amine conjugate, we employed a reverse transcriptase (RT) primer extension stop assay. The arylation of RNA has been shown to halt an RT enzyme from traversing a modification site, terminating cDNA synthesis one nucleotide prior to the modified position.^16^ We chose a 51 nt RNA with minimal secondary structure to evaluate reactivity without structural bias (Figure 3F). Briefly, the reaction was carried out by annealing the RNA with DNA-amine conjugate and treating with 2 mM cyanopyrimidine **5** overnight at 37 °C (see SI for details). After reaction, primer extension was performed and analyzed by gel electrophoresis. The results showed intense bands in close proximity to the site of nucleophilic reagent, directly adjacent to the DNA binding site (Figure 3G, H), affirming the site-selective delivery of the DNA conjugated DMAP. The most intense band corresponded to a site two nucleotides from the terminal DNA end bearing the nucleophile, which can be rationalized from the linker length of 13 atoms. To test the programmability of this strategy in modifying different sites of a desired RNA, we constructed a new DNA-DMAP conjugate complementary to a different site in the same 51nt RNA, shifted in position relative to the first case (Figure 3F, Table S1, Figure S12), and results again demonstrated site-selective reactivity at positions shifted by the new location of the DNA binding site, predominantly two nucleotides from the catalyst attachment site (Figure. 3G, Figure S13). Together, the data confirm site-localized reaction and sequence programmability of this approach.

To further test functional utility of this strategy, we then proceeded to attempt sequence-directed reactions of larger RNAs and mixed RNAs. This involves substantially increased sequence diversity, presenting a greater challenge for the site-selectivity of the targeted reaction. We first chose a 996-nt messenger RNA (mRNA) encoding green fluorescent protein (GFP) as a target, looking to evaluate success and selectivity of reaction by imaging the RNA and by measuring its gene expression. The modification of RNAs would be performed *in vitro* before transfecting them into cells to evaluate expression of the encoded proteins. We anticipated that modifying the polyA tail of the RNA might enable labeling without blocking protein translation, while any off-target modification of the protein-coding region of the mRNA would be expected to block ribosomal protein synthesis. Although aryl adducts of RNA have not yet been tested for their effects on translation, prior studies of acyl 2’-OH adducts of similar size have been shown to block ribosome progression effectively.^10^ Thus, successful modification in the polyA tail would provide a visible label on the RNA while also testing site selectivity as measured by protein expression.

For the labelling studies, we synthesized a new sulfonamide-modified pyrimidine halide reagent **SFPz** with an azide handle for labelling or pulldown applications (Figure 4A). Reactions of a short test RNA (18 nt) with this reagent in the presence of free DMAP resulted in arylated RNA bearing the azide group (Figure 4B), which could be subsequently reacted with DBCO TAMRA to yield the fluorescent conjugate, confirmed by mass spectrometry and visualized by gel electrophoresis (Figure 4C). We then employed the above strategy to visualize mRNA and its translated protein by separately labelling the polyA tails of eGFP and mCherry mRNAs with Cy5 and FAM respectively (Figures 4D,E). The strategy did not noticeably perturb the translation and consequent eGFP (green fluorescence) and mCherry (red fluorescence) expression (Figure 4E (i, ii)), over time (Figure S14). We note that both eGFP and mCherry mRNAs possess a 120 nt polyA tail, where the T20 DNA-catalyst conjugate can bind and label the RNA at multiple positions.

**Figure 4.**
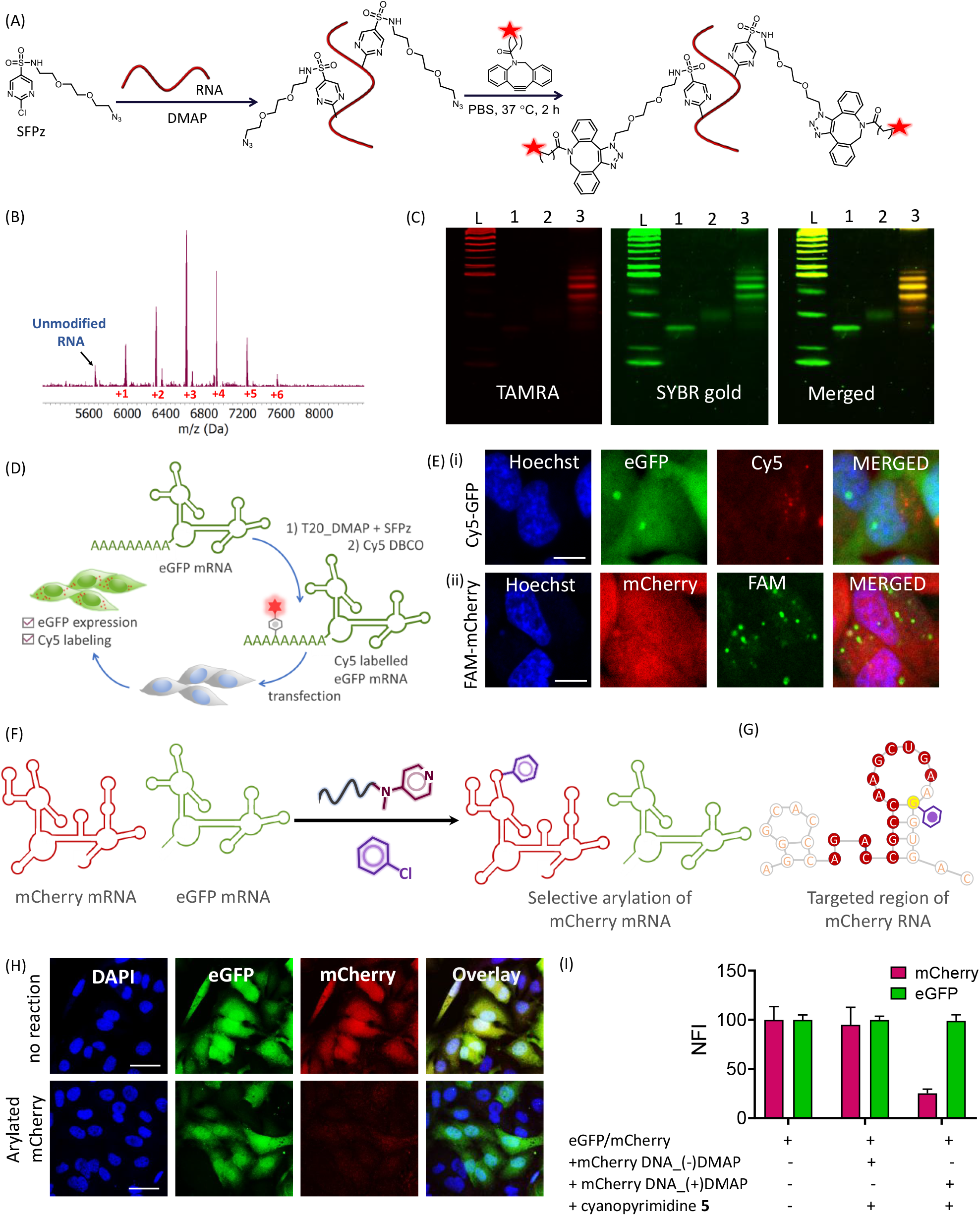
Employing sequence-directed RNA arylation in labeling and in selective control of translation. (A) Reaction scheme for synthesis of the azide-modified arylating agent **SFPz** for fluorescent labeling; (B) MALDI-TOF spectrum for reaction of 18 nt RNA with the azide-modified pyrimidine carried out with free DMAP (100 mM), confirming its catalyst responsiveness. (C) PAGE gel image of fluorescent labelling of the 18 nt RNA in (B) after click reaction with DBCO-TAMRA. Lanes: L: DNA ladder, 1: unreacted 18 nt RNA; 2: RNA modified with **SFPz**; 3: **SFPz** modified RNA labeled with TAMRA. (D) Schematic representation of reaction and subsequent transfection of polyA-labelled mRNAs. (E) Confocal microscopy images of HeLa cells transfected with (i) Cy5 labeled eGFP mRNA and (ii) FAM labeled mCherry mRNA. Scalebar: 10 μm. (F) Scheme for transcript-selective arylation; (G) Partially unpaired region of mCherry mRNA chosen for selective arylation (see Fiure S12 for full predicted structure). (H) Representative confocal images of HeLa cells after transfection with EGFP/mCherry mRNAs, showing selective knockdown of mCherry expression. Mixed mRNAs were incubated with 15mer DNA-amine conjugate in the presence of cyanopyrimidine **5** (2 mM) prior to transfection. Scalebar: 40 μm. (I) Bar graphs of green/red fluorescence ratios from cells treated with mixed RNAs after selective arylation by DNA-DMAP. Data are averages from 3 replicates; error bars show standard deviations.

Finally, to further test the sequence-specific delivery of an aryl group to a targeted mRNA in the presence of other mRNAs, we constructed a DMAP-DNA conjugate to selectively arylate a site in the protein-coding region of mCherry mRNA from a mixture of eGFP and mCherry mRNAs (Figure 4F). Since these combined RNAs constitute nearly 2000 nt of sequence, this provides a more stringent test of selectivity. Because DNA hybridization with folded RNA is well known to be sensitive to secondary structure of the target RNA,^21^ we employed folding predictions^22^ of the mCherry RNA to choose a site in the coding region likely to have a segment of unpaired structure (Figs. 4G, S15), and constructed a DNA-DMAP conjugate complementary to it (Figure S16). The 15nt DNA (Figure 4G), was designed to specifically modify mCherry with an aryl group to result in translational repression, while potentially leaving the eGFP mRNA relatively unaffected due to a lack of complementarity. We incubated a 1:1 mixture of the two mRNAs with the mCherry-complementary DNA-DMAP conjugate and cyanopyrimidine reagent **5**. After the arylation reaction, transfection of the mixture of the two mRNAs revealed a 75 % knockdown in mCherry protein expression in HeLa cells as measured by fluorescence at 590 nm (Figure 4H,I). Imaging and fluorescence analysis of the cells reveals this marked decrease in red emission signal, while green GFP fluorescence was unaffected. To rule out a simple antisense effect of the DNA on mRNA expression, we performed control experiments employing the same DNA-azide lacking the DMAP. The controls showed no decrease in the translation of either mRNA (Figure 4I). Taken together, the results provide a clear demonstration of sequence-specific arylation chemistry implemented in a mixture of messenger RNAs.

## Conclusions

Our experimental results establish a novel chemical strategy to arylate RNA site-selectively. The method employs short complementary DNA-amine conjugates to activate and direct reaction with aryl chlorides added to solution. We achieved this by identifying DMAP as both an efficient nucleophile and leaving group for aryl chloride electrophiles in aqueous solution, and by establishing that DMAP-aryl adducts are highly reactive to RNA 2’-OH groups in water, resulting in aryl ether modifications to the biopolymer. Importantly, these aryl chlorides are poorly reactive with RNA, while addition of DMAP to the solution results in a transient ammonium species that is highly reactive. Finally, to render the reaction into a site-selective modifying method, we linked DMAP to DNA oligonucleotides, employing them to deliver adducts on the RNA at specific sites programmed by the DNA sequence. We further find that the sequence-specific reaction can be carried out on biologically active mRNAs in a region that does not interfere with RNA expression, and employ it to fluorescently label the RNAs by design of a new azide-modified arylating reagent. Finally, we show that this DNA-directed arylation can be targeted in a transcript-specific way, by suppressing protein expression activity of a coding RNA while leaving the biological activity of another unaffected.

The design and translation of this technique to site-directed RNA modification is simpler and more versatile than existing methods in a number of ways. With the current DNA-mediated delivery, aryl precursors are widely available and could be synthetically modified via shorter and more economic synthetic routes than are required for previous DNA enzyme substrates,^7,8^ and does not require custom DNA synthesis nor synthetically modified nucleotides. In addition, our experiments document site-selective mRNA modification without the necessity of masking the entire RNA.^10^ Finally, while prior reports have described the use of DNA delivery of reactive groups to RNA,^13,14^ the current approach offers a robust strategy that operates under mild conditions that maintain large RNA integrity with low off-target reactivity. To the best of our knowledge, this is the first report of short complementary DNA-directed site-specific modulation and labeling of messenger RNAs.

The current results also suggest areas for future development. The current DNA conjugate design yields reaction that is not confined to a single nucleotide, but occurs over 2-3 adjacent nucleotides. While this is not a concern for most applications, it seems possible that the precision of sequence-specific arylation may be enhanced in the future by optimizing the length, position, and rigidity of the linker moiety appending the amine with the complementary DNA. Second, identifying more highly DMAP-responsive electrophiles may speed reaction, provided that they retain low RNA reactivity. Finally, it is well recognized that successful DNA hybridization to RNA depends heavily on the folding of the RNA;^22^ thus knowledge of the target RNA secondary structure is important to the current method, and testing multiple sequences and sites, as is common in the design of hybridization probes, may be necessary for optimal results. The use of DNA modifications that enhance RNA binding may also be helpful in competing with the target secondary structure.^23^

We anticipate that this strategy offers substantial opportunities for future expansion both in chemistry and in application. Addition of varied functional groups and labels to pyrimidine and triazine precursors may enable a broader range of end uses for site-directed arylation. Moreover, DMAP and other nucleophilic catalysts are well known to promote other reactivity including acyl transfer,^24^ and so we anticipate that the strategy will likely not be limited to the S_N_Ar reaction mechanism and may be translated to a broader range of RNA modification chemistries.^25^ It is also conceivable that this or related approaches might be applied directly in living cells with optimization of DNA conjugate delivery and electrophile biocompatibility. Potential future applications of this strategy may leverage site-specific modification of mRNAs to locally alter or modify structure and function for mechanistic understanding or therapeutic intervention.

## Author Contributions

E.T.K. directed the study. S.P., W.Z., S.F., and S.C. performed experiments. S.P. and E.T.K wrote the manuscript.

## Acknowledgements

We thank the U.S. National Institutes of Health (GM145357) for support.

## Conflicts of Interest

None to report.

## Entry for the Table of Contents

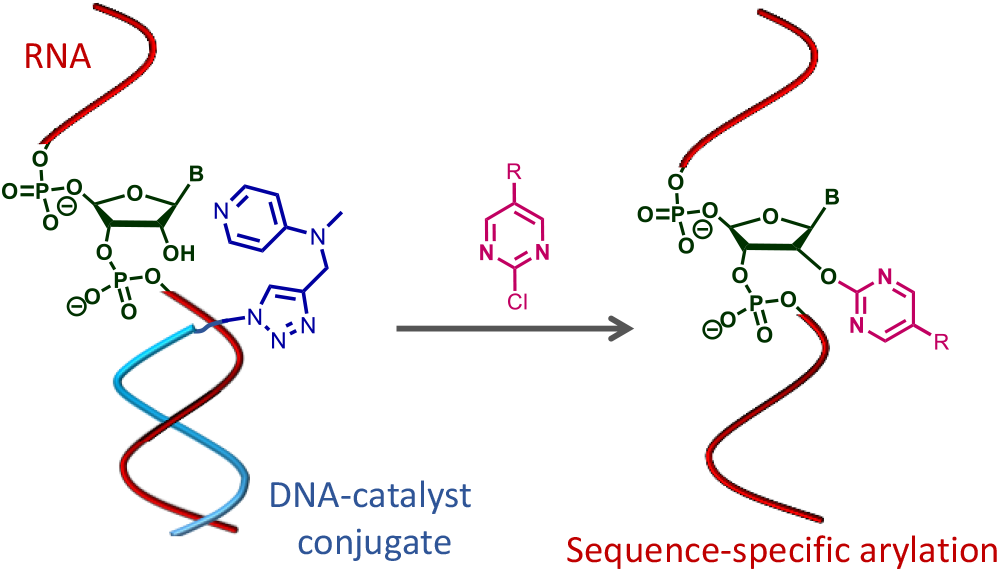

DNA oligonucleotides carrying amine nucleophilic catalysts are employed to deliver aryl groups to complementary sites in RNA under physiological conditions. The strategy can be used to sequence-specifically label protein-coding RNAs, or to knock down expression selectively, with potential applications in basic transcriptome research and in future therapies.

## Notes

### Competing Interest Statement

The authors have declared no competing interest.

