## Supporting Information for "Sequence-Specific Installation of Aryl Groups in RNA via DNA-Catalyst Conjugates"

### 1. Materials and instruments

| Reagents | Source | Identifier |
|---|---|---|
| MOPS, 1.0 M buffer, pH 7.5 | Thermo Scientific™ | # J61843.AP |
| UltraPure DNase/RNase-free Distilled water | Thermo Scientific™ | # 10977023 |
| 5M NaCl | Invitrogen™ | #AM9760G |
| 1M MgCl2 | Invitrogen™ | #AM9530G |
| SequaGel - UreaGel Concentrate | National diagnostics™ | # EC-830 |
| SequaGel - UreaGel Diluent | National diagnostics™ | #EC-840 |
| SequaGel - UreaGel Buffer | National diagnostics™ | #EC-835 |
| Ammonium Persulfate | Thermo Scientific™ | # 17874 |
| N, N, N′, N′-Tetramethyl ethylenediamine | Sigma-Aldrich™ | # 110732 |
| Methyl sulfoxide-$d_6$, for NMR | Thermo Scientific™ | #320770075 |
| 10XTBE | Thermo Scientific™ | #15581-044 |
| Orange G | Aldrich | #86,128-6 |
| Bromophenol Blue | Sigma-Aldrich™ | #114405 |
| 2-Chloro-4,6-dimethoxy-1,3,5-triazine | Sigma-AldrichTM | # 375217 |
| 2-Chloro-5-pyrimidinecarbonitrile | AA BlocksTM | # AA0020SZ |
| 2-Fluoropyrimidine | AA BlocksTM | # AA00I5WQ |
| 2-Chloropyrimidine | AA BlocksTM | # AA00336W |
| 2-Chloro-5-nitropyrimidine | AA BlocksTM | # AA003GWV |
| 2-Chloro-5-(trifluoromethyl)pyrimidine | AA BlocksTM | # AA00FC4D |
| 2-Chloro-5-methanesulfonylpyrimidine | AA BlocksTM | # AA01BFJY |
| 2-Chloropyrimidine-5-carbaldehyde | AA BlocksTM | # AA0036U8 |
| 2-Chloropyrimidine-5-carboxylic acid | AA BlocksTM | # AA00BYUB |
| 2-Chloro-4,6-pyrimidinedicarbonitrile | AA BlocksTM | # AA01HCQ2 |
| 3,6-Dichloro-1,2,4,5-tetrazine | Sigma-Aldrich™ | # 792594 |
| 4-Methylmorpholine | Sigma-Aldrich™ | # 407704 |
| Morpholine | Sigma-Aldrich™ | # 252360 |
| Imidazole | Sigma-Aldrich™ | # I2399 |
| 1-Methylimidazole | Sigma-Aldrich™ | # M50834 |
| 4-(Dimethylamino)-pyridine | Sigma-Aldrich™ | # 107700 |
| *N*-Methylpropargylamine | Sigma-Aldrich™ | # 150223 |
| Methyl sulfoxide, 99.7+%, Extra Dry, AcroSeal™ | Thermo Scientific™ | # AC326881000 |
| 96% Ethanol | Fisher Scientific™ | # BP8202-500 |
| Glycogen | Sigma-Aldrich™ | |
| 3 M sodium acetate, pH=5.2 | Thermo ScientificTM | # AM9740 |
| RNaseOUT™ Recombinant Ribonuclease Inhibitor | Invitrogen™ | # 10777019 |
| SuperScript™ II Reverse Transcriptase | Invitrogen™ | # 18064022 |
| Urea | Fisher Scientific™ | #BP169 |
| Glycerol | Sigma-Aldrich™ | #G5516 |
| 1M Tris. HCl buffer pH 7 | Invitrogen™ | #AM980G |
| dNTP mix (10 mM each) | Thermo Scientific™ | # R0194 |
| TAMRA DBCO | Click Chemistry Tools™ | # A-131 |
| RQ1 RNase-Free DNase | Promega | # M6101 |

| | | |
|---|---|---|
| Ammonium citrate dibasic | Sigma-Aldrich™ | # 09833 |
| 2′,4′,6′-Trihydroxyacetophenone monohydrate | Sigma-Aldrich™ | # T64602 |
| Hexane | Fisher Chemicals | #H292-20 |
| Ethyl acetate | Fisher Chemicals | #E145-20 |
| Dichloromethane | Fisher Chemicals | #D37-20 |
| Methanol | Fisher Chemicals | #A412-20 |
| Magnesium sulfate anhydrous | Fisher Chemicals | # M65-500 |
| DMF | Sigma-Aldrich™ | # 227056 |
| N,N-Diisopropylethylamine, 99.5+%, AcroSeal™ | Thermo Scientific™ | # 459591000 |
| N-(3-Dimethylaminopropyl)-N′-ethylcarbodiimide hydrochloride | Sigma-Aldrich™ | # E6383 |
| DMEM (1X) | Gibco | #21063-029 |
| Opti-MEM™ I Reduced Serum Medium | Gibco | # 31985-070 |
| 0.25% Trypsin-EDTA (1X) | Gibco | #25200-056 |
| Anti-Anti | Gibco | #15240-062 |
| FBS | Gibco | # A5256701 |
| Lipofectamine™ MessengerMAX™ | Invitrogen | #100027730 |
| 8-Azido-3,6-dioxaoctan-1-amine | AK Scientific | #AMTGC18997 |
| Hoechst 33 342 | Thermo Scientific | #62249 |
| SYBR gold | Invitrogen | #S11494 |
| CleanCap® EGFP mRNA | TriLink Biotechnologies | #L-7601 |
| CleanCap® mCherry mRNA (5moU) | TriLink Biotechnologies | #L-7203 |
| 2-Chloropyrimidine-5-sulfonyl chloride | AmBeed | #A179861 |
| TEMED | Sigma-Aldrich™ | #T22500 |
| TriDye™ Ultra Low Range DNA Ladder | New England BioLabs | #N0558S |

$^{1}$H NMR and $^{13}$C NMR spectra were recorded on a Bruker 400 MHz spectrometer. Chemical shifts were referenced internally to the residual solvent signal of DMSO at 2.50 ppm. Coupling constants (J) in $^{1}$H NMR are reported in Hertz (Hz), and signal multiplicities are designated as follows: s (singlet), d (doublet), t (triplet), q (quartet), m (multiplet), dd (doublet of doublets), dq (doublet of quartets), dt (doublet of triplets), and tq (triplet of quartets).

Alexa Fluor™ 488-DBCO and TAMRA-DBCO were procured from Click Chemistry Tools. RNA and DNA oligonucleotides were synthesized by IDT. SuperScript™ II Reverse Transcriptase, RNaseOUT™ Recombinant Ribonuclease Inhibitor, DNase I, and the 10 mM dNTP solution were purchased from Thermo Fisher Scientific. Oligonucleotide concentrations were determined using a NanoDrop One microvolume UV-Vis spectrophotometer. For analysis and optimization of reactions on oligonucleotides, MALDI-TOF spectrometry was employed using a Bruker MALDI Microflex LRF instrument. The samples for MALDI-TOF analysis were spotted on a MSP BigAnchor 96 BC MALDI target with a matrix solution consisting of 0.3 M trihydroxyacetophenone (in ethanol) and 0.1 M aqueous ammonium citrate (co-matrix) in the ratio 4:1 by volume. MestReNova software was used for analysis of the MALDI-TOF spectra. Fluorescence images of the PAGE gel were recorded in a Invitrogen iBright CL1500 Imaging System at $\lambda$ex = 488 nm and $\lambda$em = 520 nm for Alex488, $\lambda$ex = 633 nm and $\lambda$em =

670 nm for Cy5. Analysis and quantification of the PAGE gel images were done using ImageJ software.

**Table S1.** List of RNAs and DNAs used in this work

| **Name** | **Sequence of RNA (left to right: 5' to 3')** |
|---|---|
| | ***RNA oligo*** |
| Test 18 nt ssRNA | AUCCUGCCGACUACGCCA |
| 25 nt ssRNA | GGGACCGAAAUAAGAGAGAAAAGAA |
| 51 nt ssRNA | GGGACCGAAAUAAGAGAGAAAAGAAGAGUAAGAAGAAAUAUAA GAGCCACC |
| | ***DNA oligo*** |
| Test 18 nt ssDNA | ATCCTGCCGACTACGCCA |
| 5' end Azide DNA | /5AzideN/TCTTATTTCGGTCCC |
| Cy5 labelled primer | /5Cy5/GG TGG CTC TTA TAT T |
| Internal Azide DNA | /5AzideN/UAAGAGAGAAAAGAA |
| T20 Azide | /5AzideN/TTTTTTTTTTTTTTTTTTTT |
| mCherryDNA | /5AzideN/TCAGCTTGGCGGTCT |

**Nucleotide Sequences of mRNAs**

eGFP mRNA ORF

5'-
AUGGUGAGCAAGGGCGAGGAGCUGUUCACCGGGGUGGUGCCCAUCCUGGUCG
AGCUGGACGGCGACGUAAACGGCCACAAGUUCAGCGUGUCCGGCGAGGGCGAG
GGCGAUGCCACCUACGGCAAGCUGACCCUGAAGUUCAUCUGCACCACCGGCAA
GCUGCCCGUGCCCUGGCCCACCCUCGUGACCACCCUGACCUACGGCGUGCAGU
GCUUCAGCCGCUACCCCGACCACAUGAAGCAGCACGACUUCUUCAAGUCCGCC
AUGCCCGAAGGCUACGUCCAGGAGCGCACCAUCUUCUUCAAGGACGACGGCAA
CUACAAGACCCGCGCCGAGGUGAAGUUCGAGGGCGACACCCUGGUGAACCGCA
UCGAGCUGAAGGGCAUCGACUUCAAGGAGGACGGCAACAUCCUGGGGCACAAG
CUGGAGUACAACUACAACAGCCACAACGUCUAUAUCAUGGCCGACAAGCAGAA
GAACGGCAUCAAGGUGAACUUCAAGAUCCGCCACAACAUCGAGGACGGCAGCG
UGCAGCUCGCCGACCACUACCAGCAGAACACCCCCAUCGGCGACGGCCCCGUG
CUGCUGCCCGACAACCACUACCUGAGCACCCAGUCCGCCCUGAGCAAAGACCC

CAACGAGAAGCGCGAUCACAUGGUCCUGCUGGAGUUCGUGACCGCCGCCGGGA
UCACUCUCGGCAUGGACGAGCUGUACAAGUAA

mCherry mRNA

AGGAAAUAAGAGAGAAAAGAAGAGUAAGAAGAAAUAUAAGAGCCACCAUGGU
GAGCAAGGGCGAGGAGGACAACAUGGCCAUCAUCAAGGAGUUCAUGCGGUUC
AAGGUGCACAUGGAGGGCAGCGUGAACGGCCACGAGUUCGAGAUCGAGGGCG
AGGGCGAGGGCCGGCCCUACGAGGGCACCCAGACCGCCAAGCUGAAGGUGACC
AAGGGCGGCCCCCUGCCCUUCGCCUGGGACAUCCUGAGCCCCCAGUUCAUGUA
CGGCAGCAAGGCCUACGUGAAGCACCCCGCCGACAUCCCCGACUACCUGAAGC
UGAGCUUCCCCGAGGGCUUCAAGUGGGAGCGGGUGAUGAACUUCGAGGACGG
CGGCGUGGUGACCGUGACCCAGGACAGCAGCCUGCAGGACGGCGAGUUCAUCU
ACAAGGUGAAGCUGCGGGGCACCAACUUCCCCAGCGACGGCCCCGUGAUGCAG
AAGAAGACCAUGGGCUGGGAGGCCAGCAGCGAGCGGAUGUACCCCGAGGACGG
CGCCCUGAAGGGCGAGAUCAAGCAGCGGCUGAAGCUGAAGGACGGCGGCCACU
ACGACGCCGAGGUGAAGACCACCUACAAGGCCAAGAAGCCCGUGCAGCUGCCC
GGCGCCUACAACGUGAACAUCAAGCUGGACAUCACCAGCCACAACGAGGACUA
CACCAUCGUGGAGCAGUACGAGCGGGCCGAGGGCCGGCACAGCACCGGCGGCA
UGGACGAGCUGUACAAGAGCGGCAACUGAGCGGCCGCUUAAUUAAGCUGCCUU
CUGCGGGGCUUGCCUUCUGGCCAUGCCCUUCUUCUCUCCCUUGCACCUGUACC
UCUUGGUCUUUGAAUAAAGCCUGAGUAGGAAGAAAAAAAAAAAAAAAAAAAA
AAAAAAAAAAAAAAAAAAAAAAAAAAAAAAAAAAAAAAAAAAAAAAAAAAAAAA
AAAAAAAAAAAAAAAAAAAAAAAAAAAAAAAAAAAAAAAAAAAAAAAAAA

### 2. Experimental Procedures

#### RNA reaction protocol with arylating reagents

All reactions for the arylation of RNA were conducted in sterile 200 µL PCR tubes. 4.7 µL of 20 µM RNA was taken, and 3.3 µL of SHAPE 3.3X Buffer (consisting of 333 mM MOPS pH 7.5, 333 mM NaCl, and 20 mM $MgCl_2$) in water was added. Freshly prepared stocks of aryl halides and amine nucleophiles in DMSO/water were added to the reaction mixture to a final concentration of 100 mM. The reactions were incubated for 18 h or desired time at 37 °C, and subsequently purified by precipitation and washing with ethanol. For investigation of reactivity with different aryl halides, parallel reactions of the RNA in absence of DMAP, and reactivity with identical DNA sequence in presence of DMAP were measured. The level of arylation were measured by MALDI-TOF and analyzed using MestReNova software.

#### Isolation of RNA by ethanol precipitation

To the 10 µL reaction mixture of RNA taken in a 1.5 mL sterile centrifuge tube, 10 uL of 3 M NaOAc (pH 5.2), and 1 µL 20 mg/mL glycogen was added, followed by 89 µL of nuclease-free water. The solution was mixed well before adding 500 µL of ice-cold ethanol (96%). The mixture was vortexed for 30 s and stored at -80 °C for 2 h. Next the mixture was centrifuged at 14k RPM for 20 min at 4 °C. The pellet formed was washed with 70 % ethanol (twice) and dried in air. The obtained pellet was stored at -80 °C for future use or dissolved in aqueous buffer for further experiments. Concentration of the isolated RNA was determined using a Nanodrop instrument.

**Click reaction of DNA azides with propargyl-linked amine moieties**

Conjugation of the propargyl amine with the DNA azide was performed using Cu(I) catalyzed click reaction. CuBr solution 1 mg/mL was prepared in a solution containing tBuOH, DMSO and water. In a 100 µL solution of azide-modified DNA (20 µM), 40 µL of 4 mM propargylated amine nucleophile solution in aqueous buffer with minimum DMSO, and 100 µL of CuBr solution was added. The reaction mixture was incubated at 37 °C for 18 h for complete reaction. The modified DNA was isolated by ethanol precipitation and characterixed by MALDI-TOF mass spectrometry.

**Site-specific RNA arylation**

10 µM of 25 nt ssRNA was mixed with 12 µM DMAP-conjugated DNA in 20 mM Tris.HCl, 50 mM NaCl, pH 7.4, and annealed at 70 °C for 5 min and cooled slowly to 4 °C. The desired concentration of cyanopyrimidine **5** or other arylating agent was added, and 3 µL MOPS buffer was added to a final concentration of 100 mM, pH 7.5 containing 100 mM NaCl, 6 mM $MgCl_2$. The reaction mixtures were incubated at 37 °C for 18 h. Next, the reaction mixture was treated with DNase (3 µL) for 1 h, and an EDTA solution (1.5 µL) was added before heating at 65 °C for 5 min to inactivate the DNase. Finally, the RNA was isolated by ethanol precipitation as described above.

**PAGE analysis of reverse transcriptase (RT) stops**

The RT stop assay was performed as described previously with some modifications.[1] The **51 nt ssRNA** (sequence provide above) was used as the template and the 15 nt long primer denoted as **Cy5 labeled primer** has 5' Cy5 label to help visualize the cDNA products after reverse transcription. The cyanopyrimidine **5** treated RNA was annealed to 65 °C with primer and dNTP mix (10 mM) in 20 mM Tris. HCl, 50 mM NaCl, pH 7.5; and cooled slowly to 4 °C. For reverse transcription reaction, 3 µL 5X First-Strand Buffer, 1 µL DTT (0.1 M), 0.4 µL RNaseOUT, 0.27 µL SuperScript II (200 U/µL) and nuclease-free water were sequentially added to make up the final volume of 10 µL. The reaction was incubated at 25 °C, and 42 °C for 2 h on a thermocycler PCR instrument. After the reaction, equal volume (10 µL) loading dye (8 M Urea, 0.05% Orange G, 0.05% Bromophenol blue in water) was added, mixed well, and denatured at 95 °C for 5 min, and loaded on a denaturing 20 % polyacrylamide gel and run at 25 mA in 1 x TBE. The cDNA products were visualized using Invitrogen iBright CL1500 Imaging System and Cy5 laser, and quantified using ImageJ. All experiments were performed in triplicates.

**Selective Arylation and Transfection of eGFP and mCherry mRNA**

0.9 µg of DMAP attached mCherryDNA was dissolved in 4 µL Tris. HCl (20 mM, pH 7.4, NaCl 50 mM). mCherryDNA-DMAP conjugate was added to the 0.85 µg of mCherry and eGFP mRNA and briefly annealed at 60 °C for 3 min and slowly cooled to room temperature and chilled on ice for 5 min. Then 3 µL MOPS buffer (composition mentioned above) was added. Next, from a 20 mM solution of cyanopyrimidine **5** prepared in MOPS buffer containing minimal DMSO, 1 µL was added to the reaction, followed by brief vortexing and centrifugation. The reaction mixture was then incubated at 30 °C overnight.

After 18 h, DNAse (4 µL) was added, with 1.5 uL of DNAse buffer, and incubated at 37 °C for 2 h. Then DNAse stop solution was added, and the RNA was isolated using Zymo clean and concentrator kit using manufacturer's protocol. The concentration of the isolated RNA was measured using a Nano drop instrument.

HeLa and HEK293T cells were obtained from the American Type Culture Collection (ATCC). HeLa cells were cultured in Dulbecco's Modified Eagle's medium (DMEM) supplemented with 10 % fetal bovine serum (FBS) and 1% anti-anti solution at 37 °C in a humidified atmosphere containing 5% $CO_2$. The transfection experiment was designed in a 96 well plate with a transparent flat bottom for fluorescence measurement. 200 ng RNA was transfected in each well using Lipofectamine MessengerMax per the manufacturer's instructions. The protein expression was monitored over various time points (6 h to 48 h), measuring the fluorescence emission corresponding to eGFP and mCherry in a microplate reader.

For fluorescence imaging of the selective blocking of translation, HeLa cells were seeded in confocal dish, and similar experimental procedure was followed. After 24 h, nuclear staining dye Hoechst (200 nM) was added to the solution and incubated for 20 min. Then the cells were washed with PBS, and fresh media was added. The eGFP, mCherry, and Hoechst emission were imaged with a laser confocal fluorescence microscope.

**Reaction of TAMRA DBCO with SFPz**

18 nt ssRNA (10 µL, 20 µM) was arylated using **SFPz** (100 mM) in presence of DMAP in MOPS buffer. After reaction, the RNA was isolated using ethanol precipitation and further washed with 70 % ethanol, and air dried. The dried pellet was dissolved in 4 µL nuclease free water. Then 200 µM of TAMRA DBCO or other relevant dye was added to this arylated RNA solution. Then 20 µL PBS (1X) pH 7.4 was further added and the reaction was incubated at 35 °C for 2 h. For isolation of short RNA, 11 µL sodium acetate was added, followed by 1 µL glycogen (20 mg/mL) and 60 uL water. Then 800 µL ethanol (96 %) was added, and the reaction mixture was mixed well and incubated at -80 °C for 1h. Then the solution was centrifuged at 14k RPM for 20 min at 4 °C, and the obtained pellet was further washed with 70 % ethanol twice. The pellet was dried in air for 10-15 min.

**Site-specific arylation of poly A tails in eGFP and mCherry mRNA**

0.7 µg of eGFP mRNA was dissolved in a buffer containing 20 mM Tris.HCl, pH 7.5, containing 50 mM NaCl. 0.72 µg of 15 nt DNA-DMAP oligo complementary to eGFP 3' UTR region was added with 4 µL of the same buffer. The solution was annealed at 65 °C for 5 min and cooled slowly to rt over 2 h. Then SFPz was added to a final concentration of 2 mM. The reaction was incubated at 30 °C for 24 h. The reaction mixture was treated with 100 µM

TAMRA DBCO solution, at 35 °C for 2 h and RNA was isolated using Zymo RNA Clean and Concentrator kit using manufacturer's protocol.

**Electrophoretic mobility shift assays**

8 µL of 18 nt ssRNA (4 µM) and modified RNAs were prepared in Tris. HCl, 50 mM NaCl. Next, 10 µL of loading dye was added to each sample. 20 % denaturing urea PAGE gel was cast. RNA and modified RNA samples were loaded after mixing with loading dye. 5 µL of DNA ladder was used as a reference. The gel was run for 1 h at current 25 mA. Then the gel was stained in SYBR gold solution for 10 min and destained for 5 min before imaging using the following channels: ex, 515, em 585-615, and ex 485, em 545-585nm.

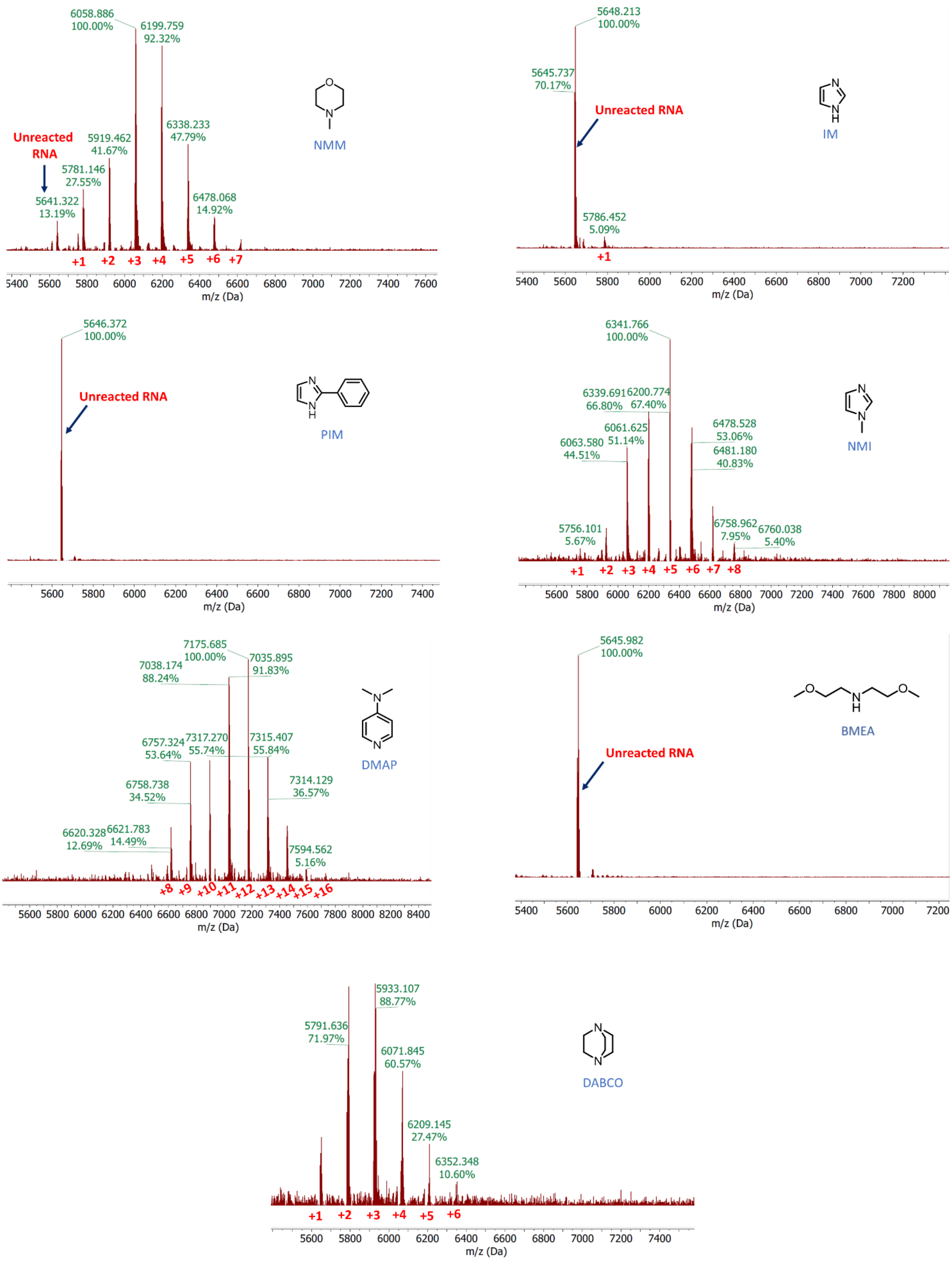


**Figure S1**. MALDI-TOF mass spectra documenting yields of reactions with 18 nt RNA using aryl chlorides with amine nucleophiles. Conditions for the screening: MOPS buffer 200 mM, pH 7.5, 100 mM NaCl, 6 mM $MgCl_2$, 37 °C, 10 μM RNA, 100 mM amine nucleophiles, 100 mM aryl chloride **11**. Red numerals indicate number of adducts.

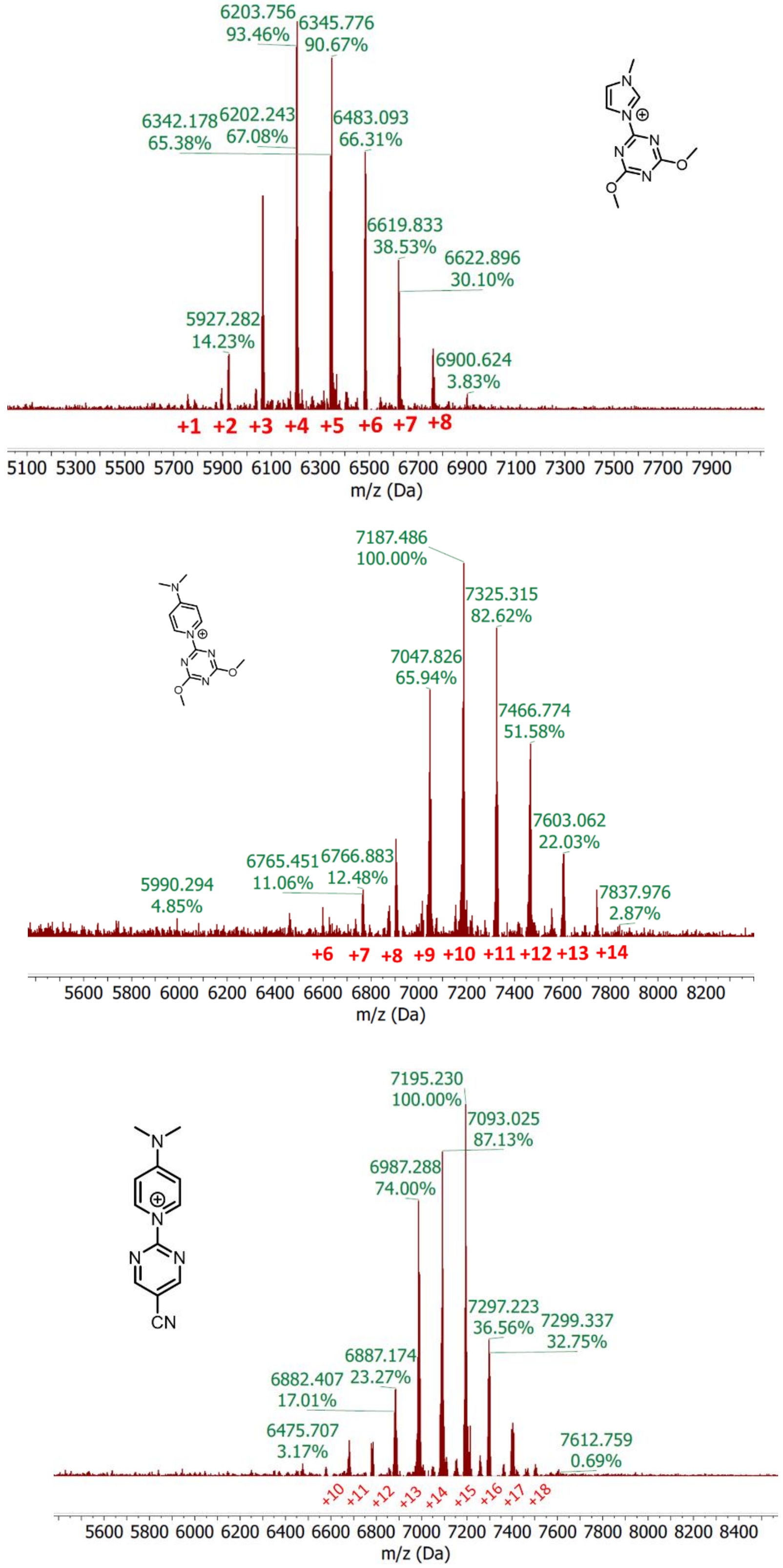


**Figure S2**. Representative MALDI-TOF mass spectra of 18 nt RNA treated with preformed complex of amine nucleophile DMAP with the electrophiles shown. Conditions for the reactions: MOPS buffer 200 mM, pH 7.5, 100 mM NaCl, 6 mM $MgCl_2$, 37 °C, 10 μM RNA, 100 mM aryl-ammonium conjugate.

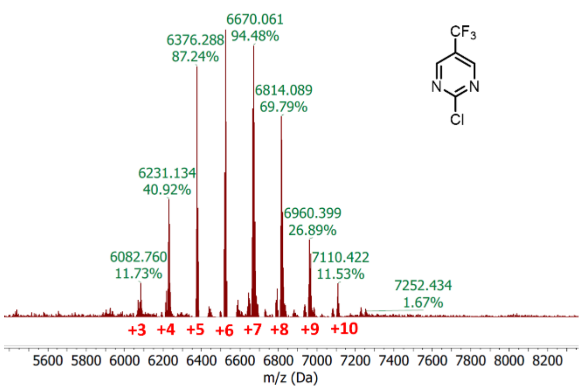
6670.061
94.48%
6376.288
87.24%
6814.089
69.79%
6231.134
40.92%
6960.399
26.89%
6082.760
11.73%
7110.422
11.53%
7252.434
1.67%
$CF_3$
N
N
Cl
+3 +4 +5 +6 +7 +8 +9 +10
5600 5800 6000 6200 6400 6600 6800 7000 7200 7400 7600 7800 8000 8200
m/z (Da)

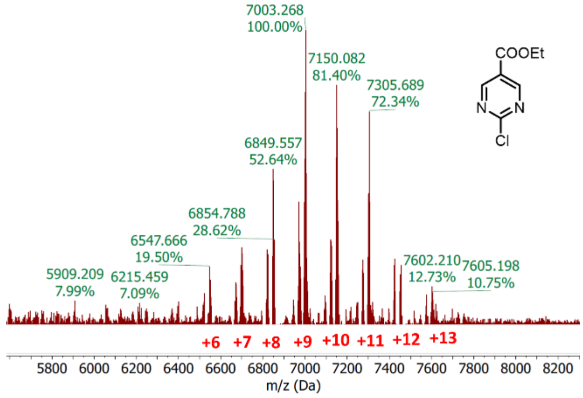
7003.268
100.00%
7150.082
81.40%
7305.689
72.34%
6849.557
52.64%
6854.788
28.62%
6547.666
19.50%
5909.209
7.99%
6215.459
7.09%
7602.210
12.73%
7605.198
10.75%
COOEt
N
N
Cl
+6 +7 +8 +9 +10 +11 +12 +13
5800 6000 6200 6400 6600 6800 7000 7200 7400 7600 7800 8000 8200
m/z (Da)

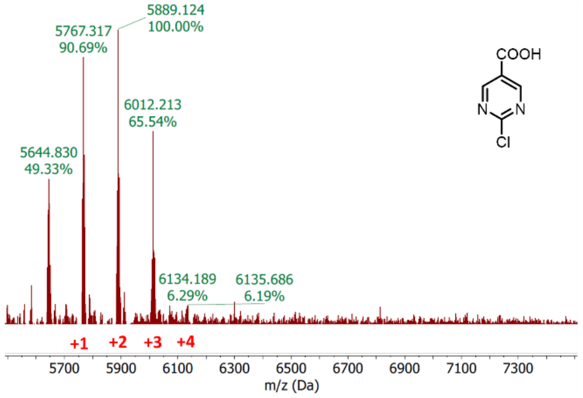
5889.124
100.00%
5767.317
90.69%
6012.213
65.54%
5644.830
49.33%
6134.189
6.29%
6135.686
6.19%
COOH
N
N
Cl
+1 +2 +3 +4
5700 5900 6100 6300 6500 6700 6900 7100 7300
m/z (Da)

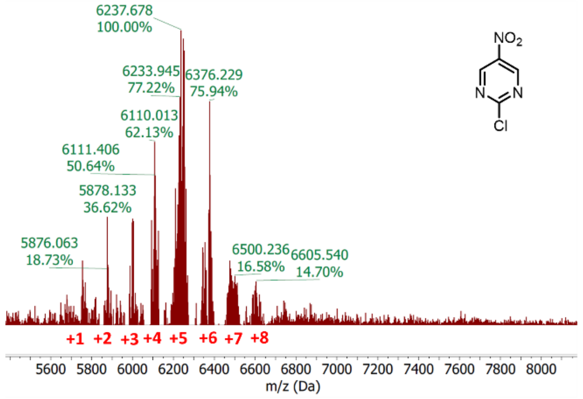
6237.678
100.00%
6233.945
77.22%
6376.229
75.94%
6110.013
62.13%
6111.406
50.64%
5878.133
36.62%
5876.063
18.73%
6500.236
16.58%
6605.540
14.70%
$NO_2$
N
N
Cl
+1 +2 +3 +4 +5 +6 +7 +8
5600 5800 6000 6200 6400 6600 6800 7000 7200 7400 7600 7800 8000
m/z (Da)

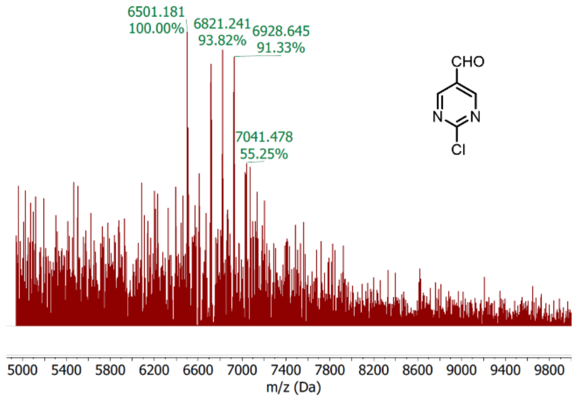
6501.181
100.00%
6821.241
93.82%
6928.645
91.33%
7041.478
55.25%
CHO
N
N
Cl
5000 5400 5800 6200 6600 7000 7400 7800 8200 8600 9000 9400 9800
m/z (Da)

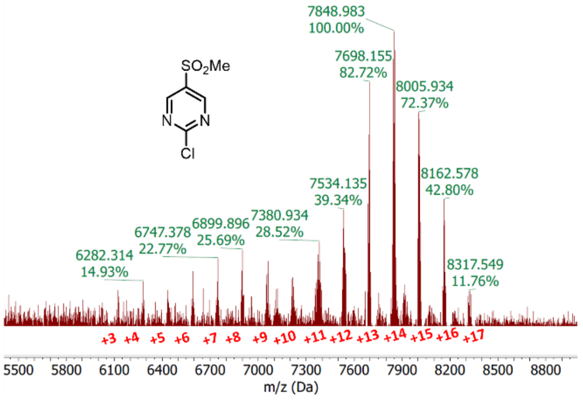
7848.983
100.00%
7698.155
82.72%
8005.934
72.37%
8162.578
42.80%
7534.135
39.34%
7380.934
28.52%
6899.896
25.69%
6747.378
22.77%
6282.314
14.93%
8317.549
11.76%
$SO_2Me$
N
N
Cl
+3 +4 +5 +6 +7 +8 +9 +10 +11 +12 +13 +14 +15 +16 +17
5500 5800 6100 6400 6700 7000 7300 7600 7900 8200 8500 8800
m/z (Da)

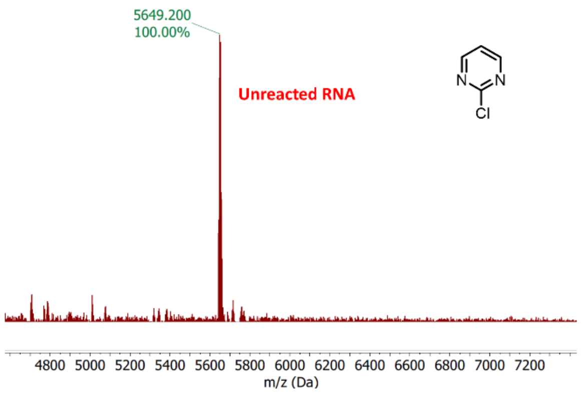
5649.200
100.00%
Unreacted RNA
N
N
Cl
4800 5000 5200 5400 5600 5800 6000 6200 6400 6600 6800 7000 7200
m/z (Da)

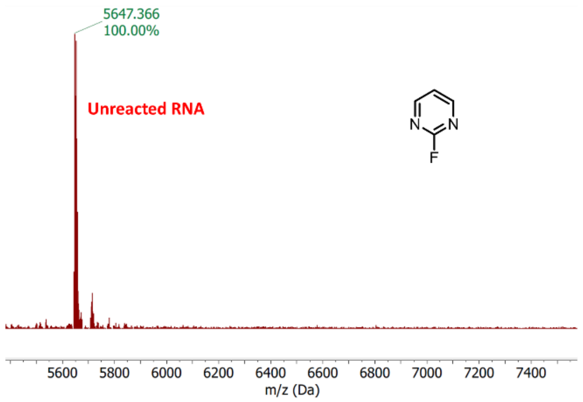
5647.366
100.00%
Unreacted RNA
N
N
F
5600 5800 6000 6200 6400 6600 6800 7000 7200 7400
m/z (Da)

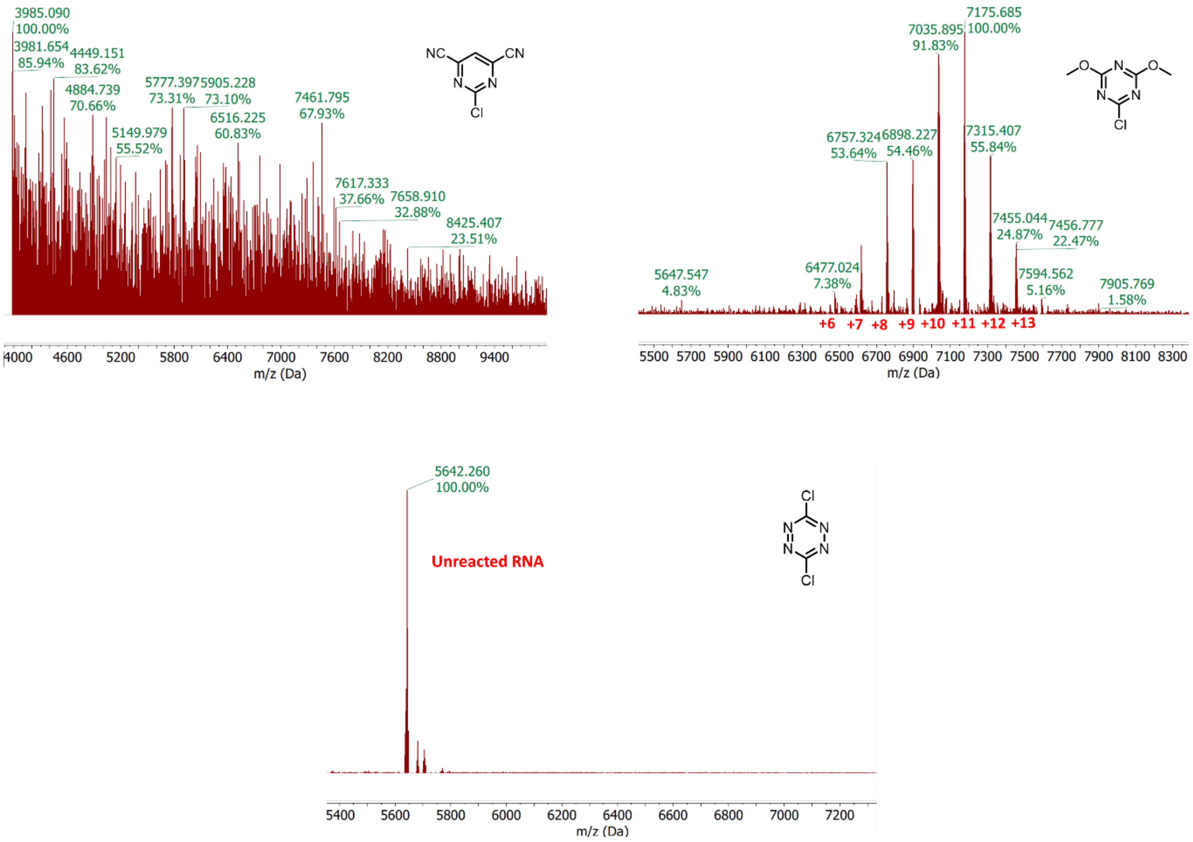


**Figure S3**. Representative MALDI-TOF mass spectra of 18 nt RNA treated with DMAP and varied aryl halides. Conditions: 100 mM DMAP, 100 mM aryl halide in MOPS buffer (100 mM, pH 7.5; 100 mM NaCl, 6 mM $MgCl_2$) for 18 h at 37 °C.

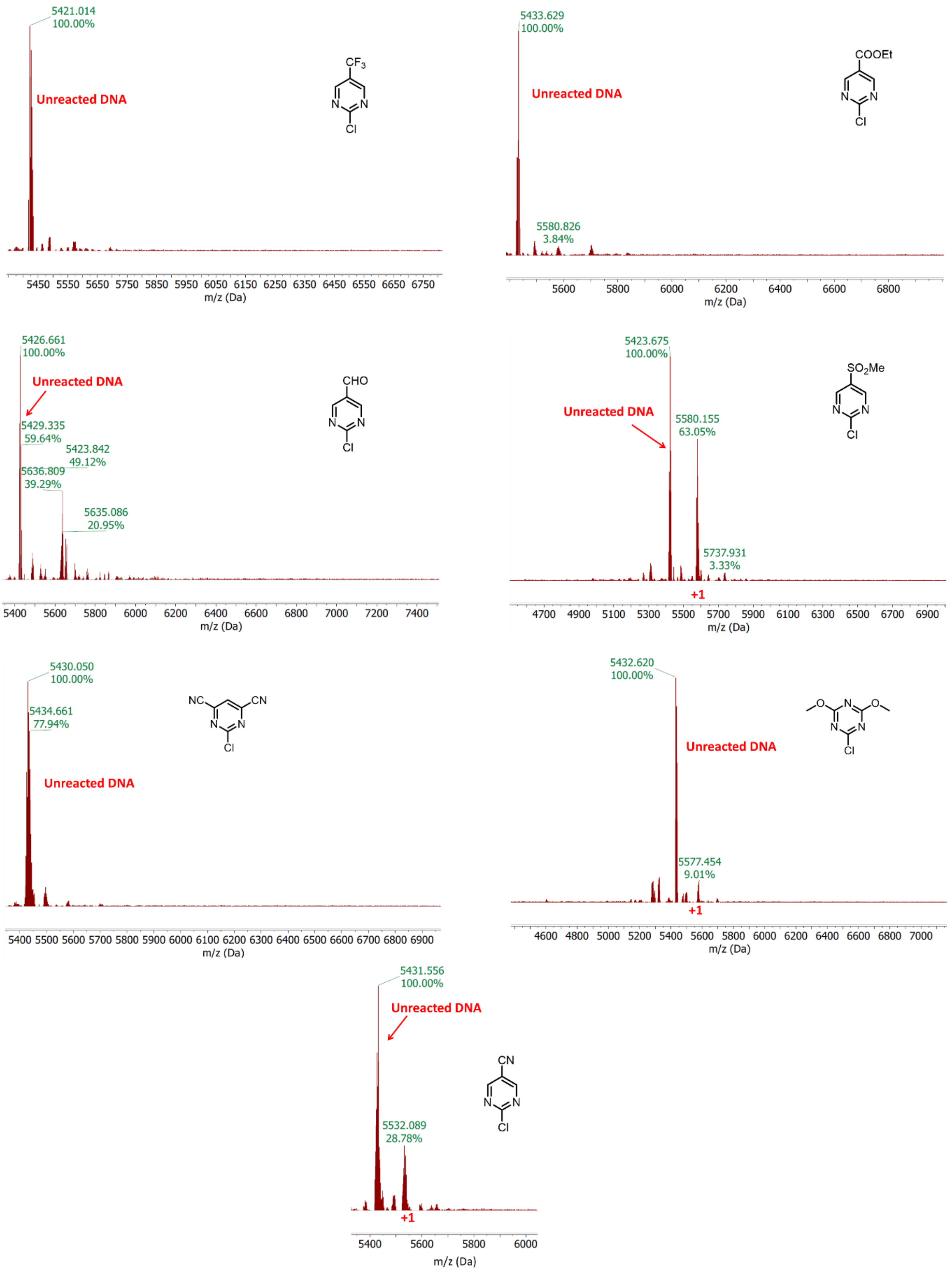


**Figure S4**. Representative MALDI-TOF mass spectra of control 18 nt DNA treated with DMAP and varied aryl halides. DNA is identical in sequence to RNA in Fig. S2; low reaction yields for DNA under conditions that provide substantial yields with RNA provide evidence of selectivity for the 2′-OH group of RNA. Conditions were the same as in Fig. S2.

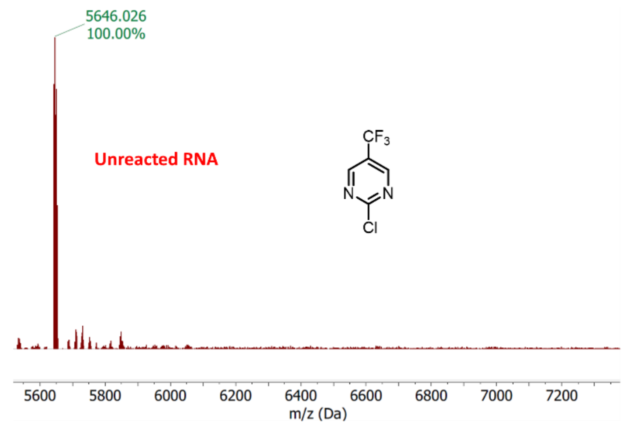
5646.026
100.00%
Unreacted RNA
$CF_3$
N
N
Cl
5600 5800 6000 6200 6400 6600 6800 7000 7200
m/z (Da)

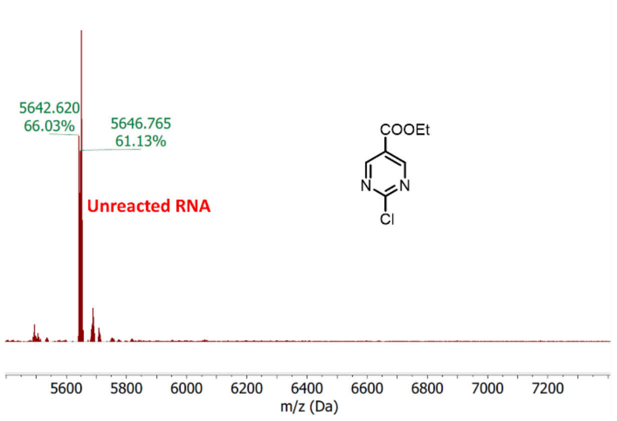
5642.620
66.03%
5646.765
61.13%
Unreacted RNA
COOEt
N
N
Cl
5600 5800 6000 6200 6400 6600 6800 7000 7200
m/z (Da)

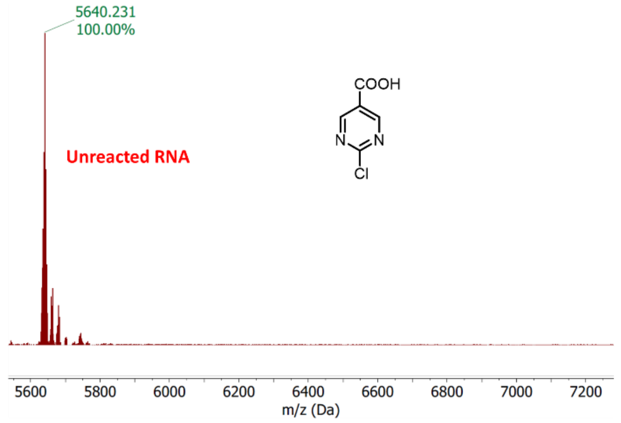
5640.231
100.00%
Unreacted RNA
COOH
N
N
Cl
5600 5800 6000 6200 6400 6600 6800 7000 7200
m/z (Da)

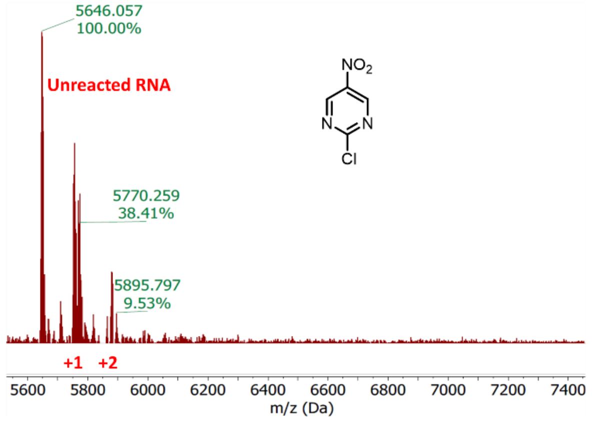
5646.057
100.00%
Unreacted RNA
$NO_2$
N
N
Cl
5770.259
38.41%
5895.797
9.53%
+1
+2
5600 5800 6000 6200 6400 6600 6800 7000 7200 7400
m/z (Da)

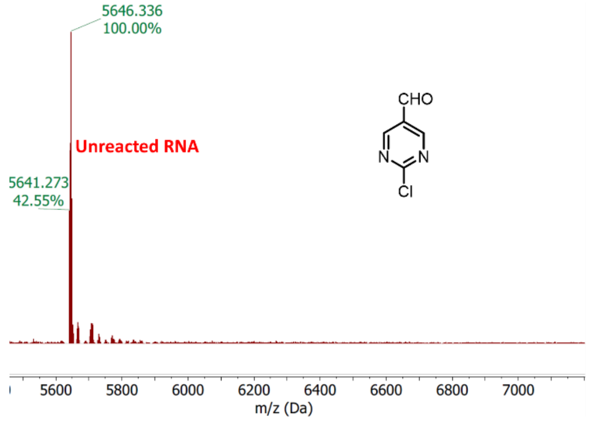
5646.336
100.00%
Unreacted RNA
5641.273
42.55%
CHO
N
N
Cl
5600 5800 6000 6200 6400 6600 6800 7000
m/z (Da)

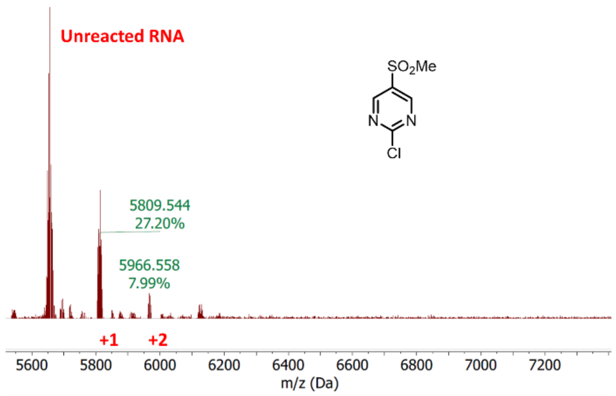
Unreacted RNA
$SO_2Me$
N
N
Cl
5809.544
27.20%
5966.558
7.99%
+1
+2
5600 5800 6000 6200 6400 6600 6800 7000 7200
m/z (Da)

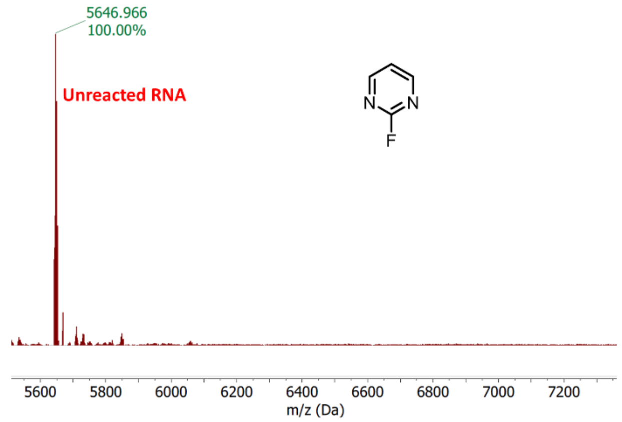
5646.966
100.00%
Unreacted RNA
N
N
F
5600 5800 6000 6200 6400 6600 6800 7000 7200
m/z (Da)

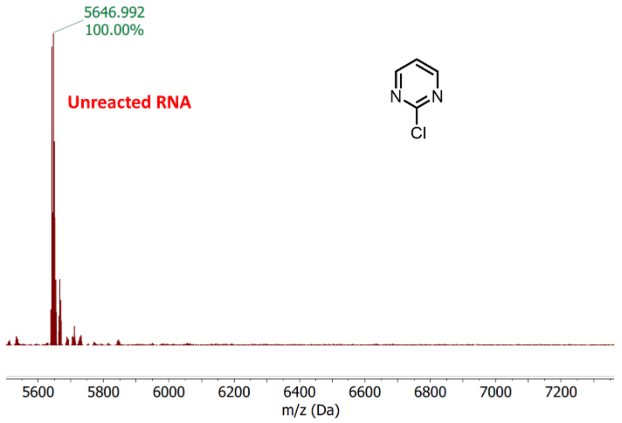
5646.992
100.00%
Unreacted RNA
N
N
Cl
5600 5800 6000 6200 6400 6600 6800 7000 7200
m/z (Da)

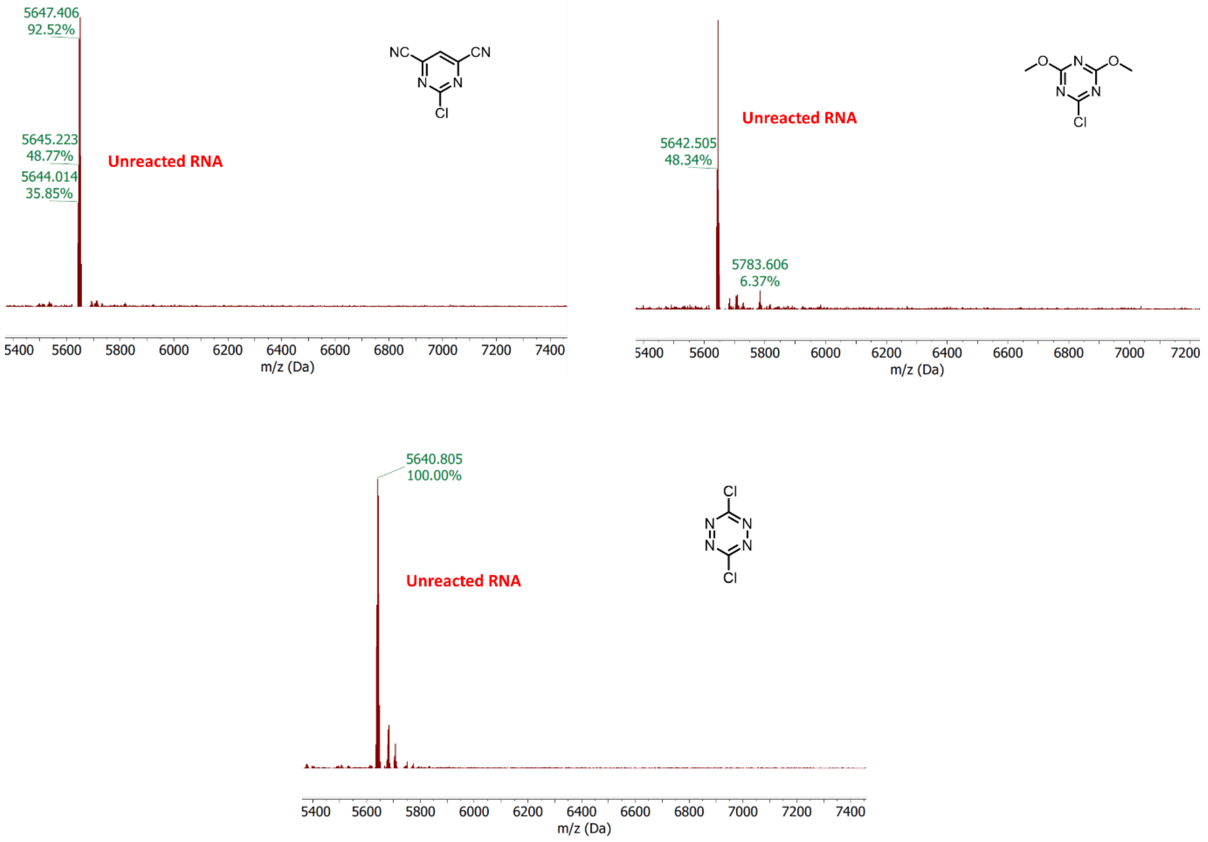


**Figure S5**. Representative MALDI-TOF mass spectra of 18 nt RNA treated with varied aryl halides in absence of amine nucleophile, DMAP. Reactions conditions were identical to that in Figure S2; low reaction yields for RNA in the absence of DMAP provide evidence for the essential role of DMAP in promoting the SNAR reactions with RNA.

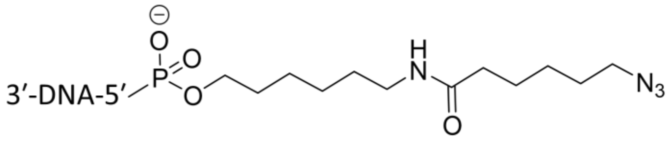


**Figure S6**. Structure of the azide linker used in this study.

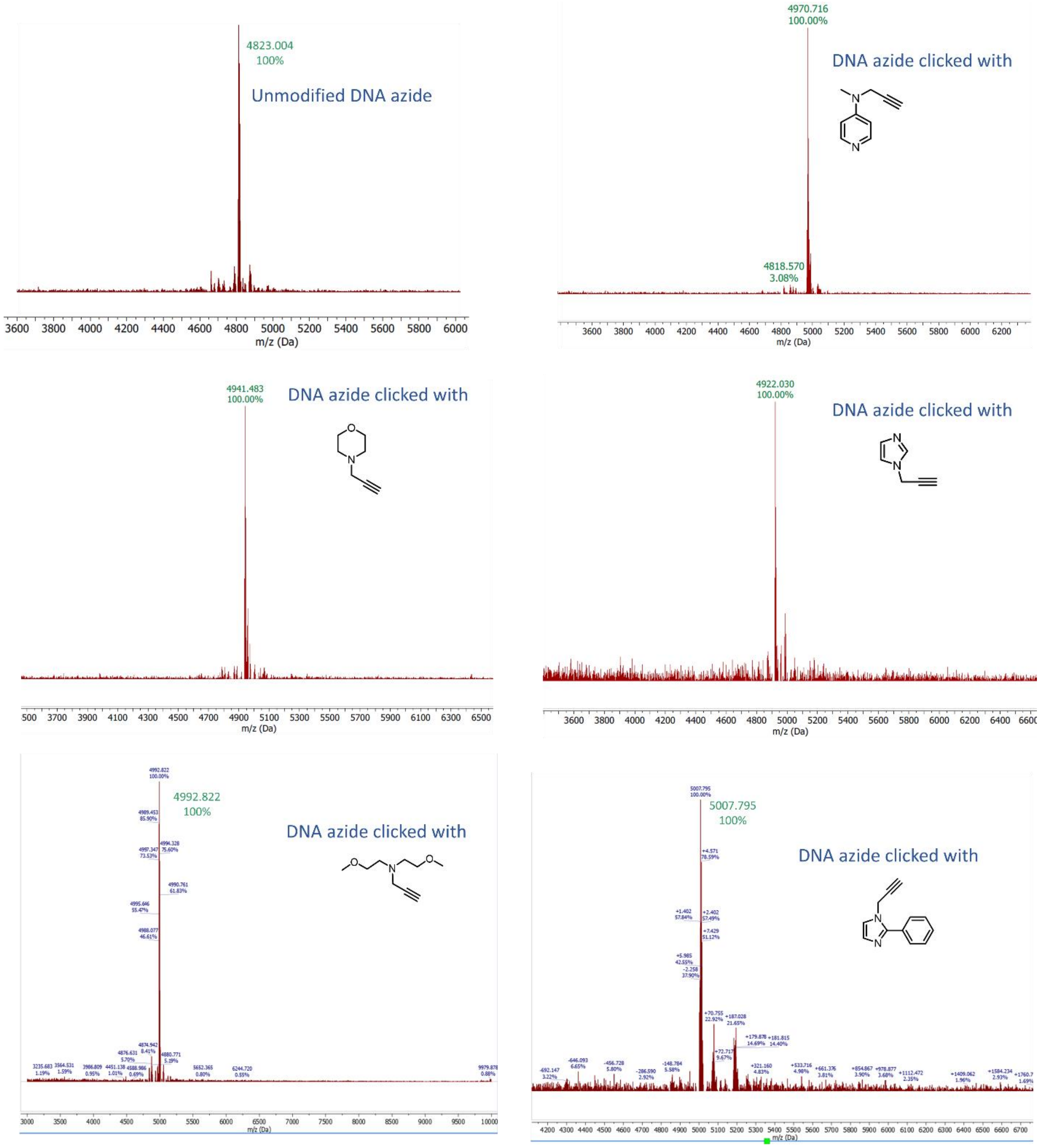


**Figure S7**. MALDI-TOF mass spectra of DNA oligonucleotide (5′ terminal DNA-Azide) modified with three different amines, confirming high yields of the expected conjugates.

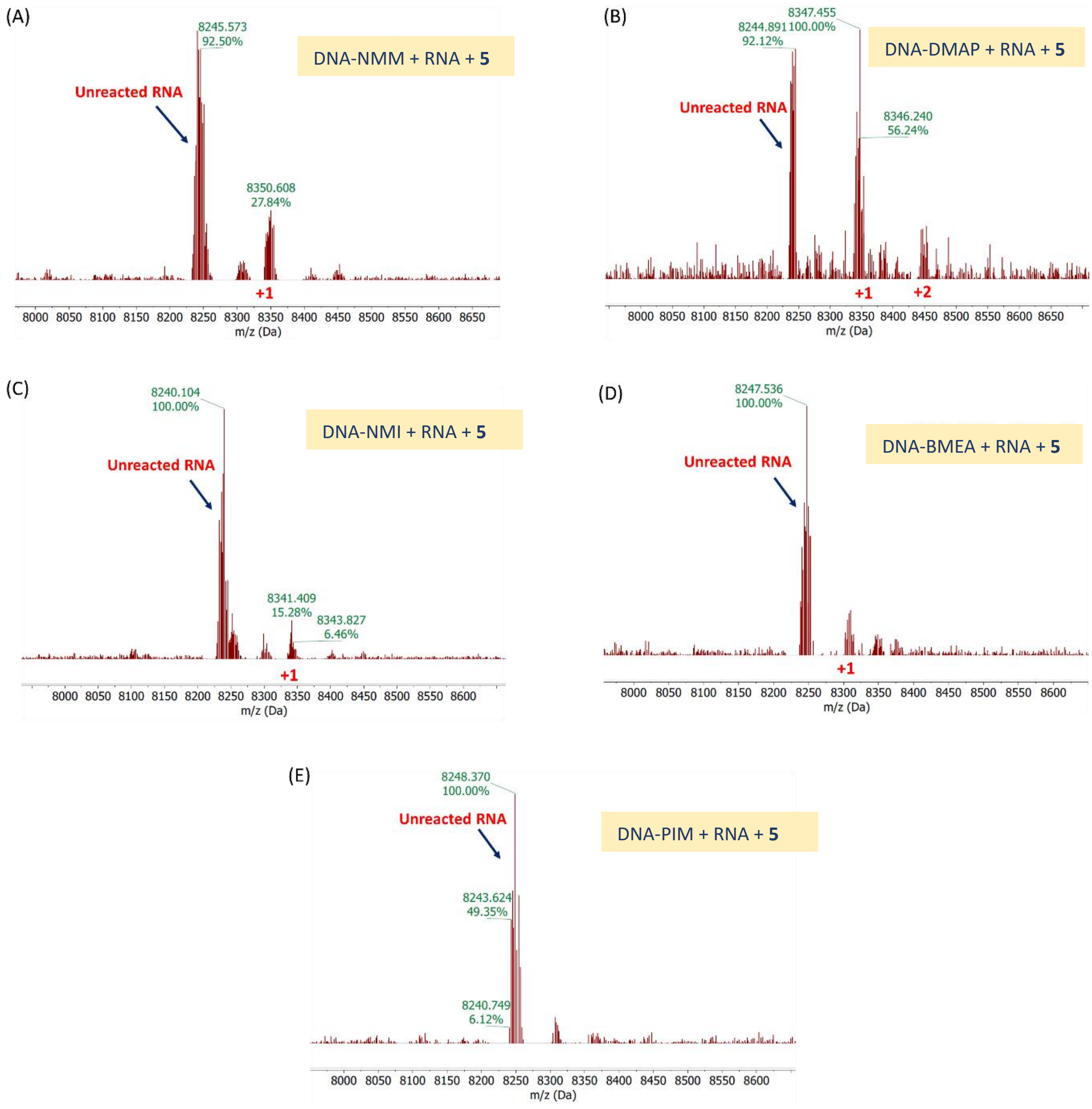


**Figure S8**. MALDI-TOF mass spectra of 25 nt RNA after DNA-directed modification with pyrimidine electrophile **5**. Spectrum of RNA in reactions with DNA-amine (A) NMM, (B) DMAP, (C) NMI, (D) BMEA, and (E) PIM. Reaction conditions: 1 mM electrophile **5**, MOPS buffer (100 mM, pH 7.5; 100 mM NaCl, 6 mM $MgCl_2$) for 18 h at 37 °C. The results confirm highest reactivity in the presence of the DMAP conjugate (B).

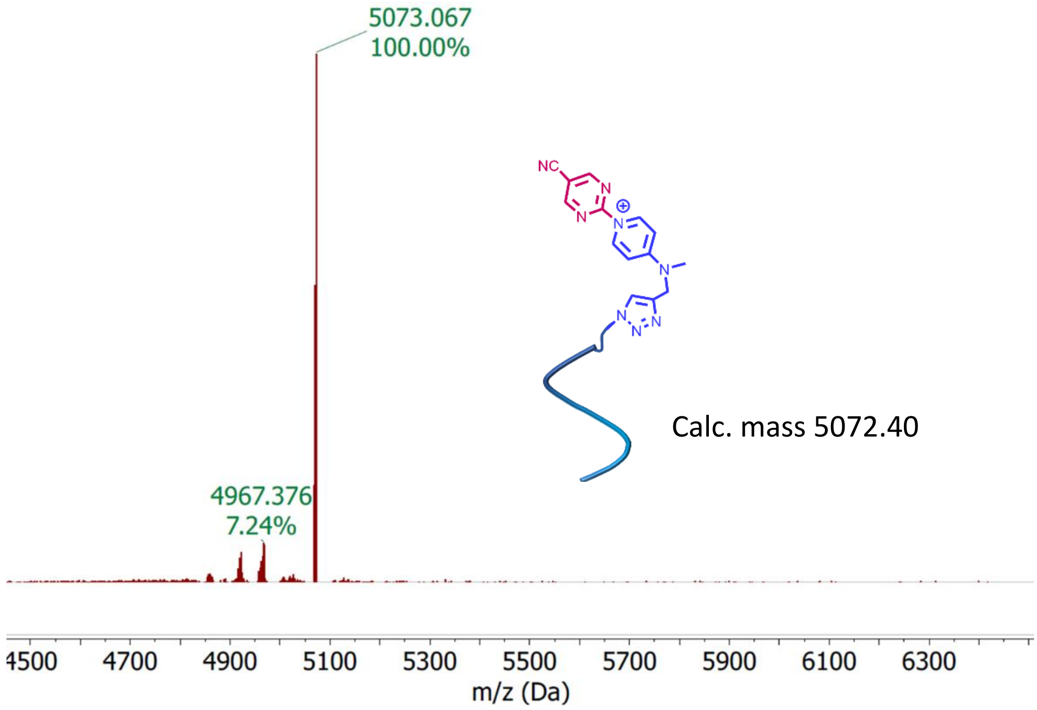


**Figure S9**. MALDI-TOF mass spectrum of DNA-DMAP conjugate after incubation with cyanopyrimidine **5**, confirming the mass of the ammonium reactive intermediate proposed to form *in situ*.

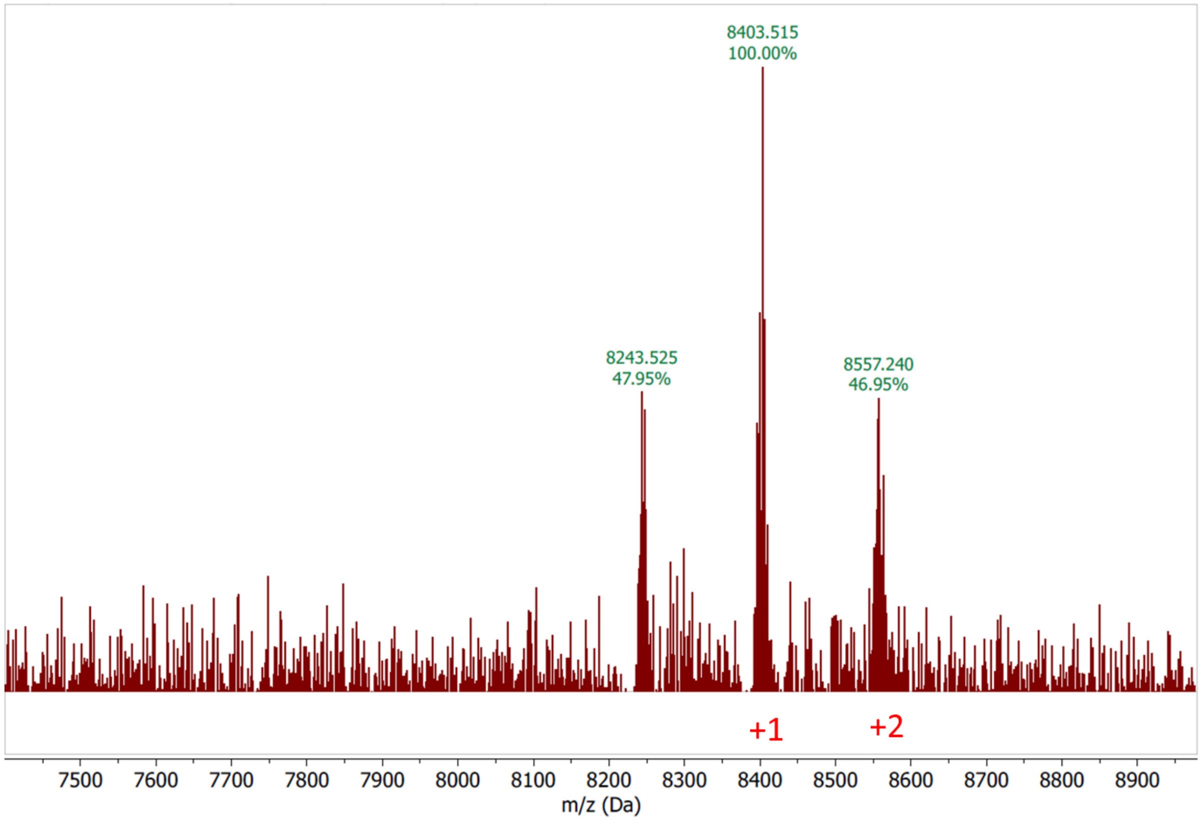


**Figure S10**. MALDI-TOF mass spectra of 25 nt RNA after DNA-directed modification with pyrimidine electrophile **7**.

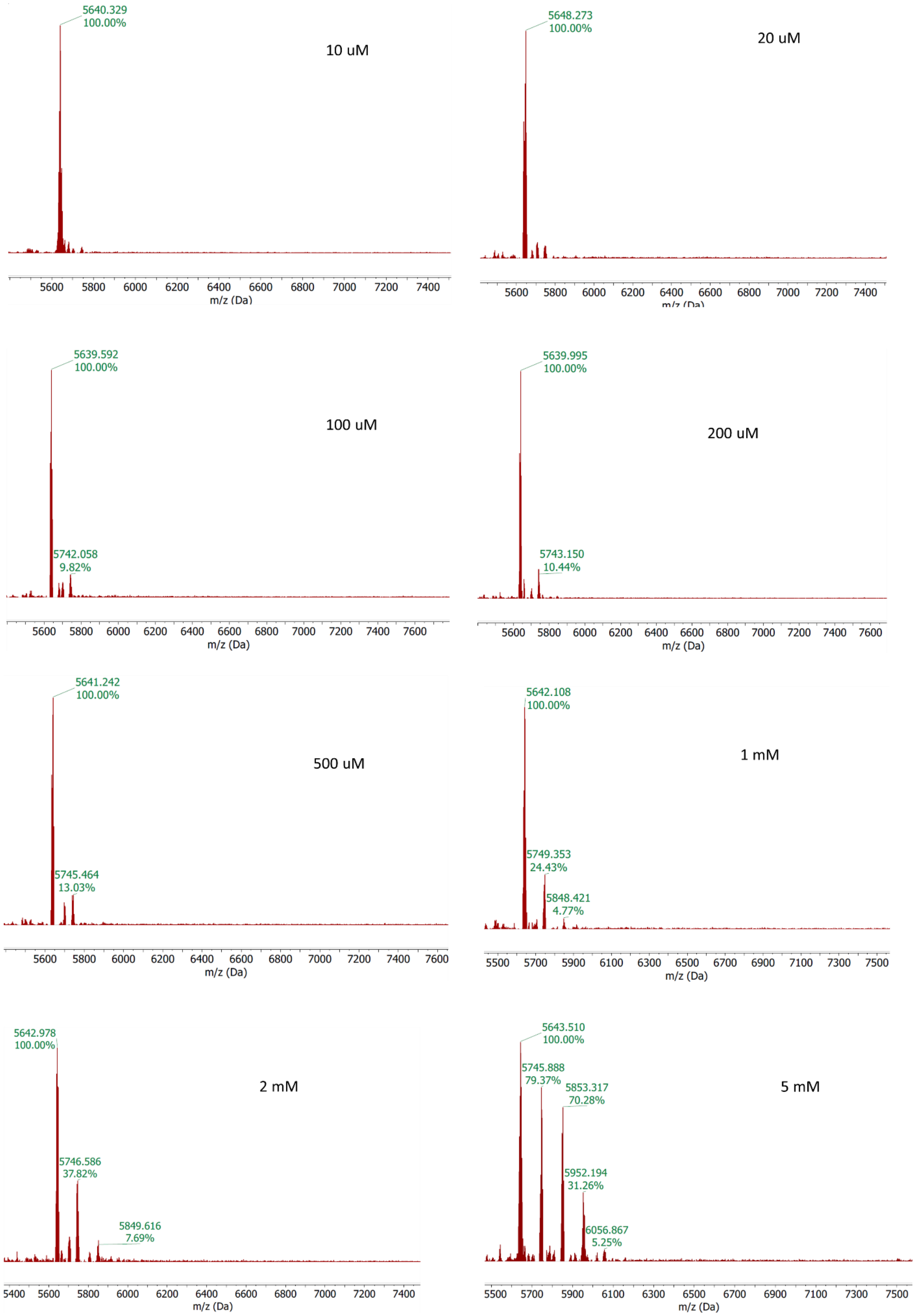
5640.329
100.00%
10 uM
m/z (Da)
5648.273
100.00%
20 uM
m/z (Da)
5639.592
100.00%
5742.058
9.82%
100 uM
m/z (Da)
5639.995
100.00%
5743.150
10.44%
200 uM
m/z (Da)
5641.242
100.00%
5745.464
13.03%
500 uM
m/z (Da)
5642.108
100.00%
5749.353
24.43%
5848.421
4.77%
1 mM
m/z (Da)
5642.978
100.00%
5746.586
37.82%
5849.616
7.69%
2 mM
m/z (Da)
5643.510
100.00%
5745.888
79.37%
5853.317
70.28%
5952.194
31.26%
6056.867
5.25%
5 mM
m/z (Da)

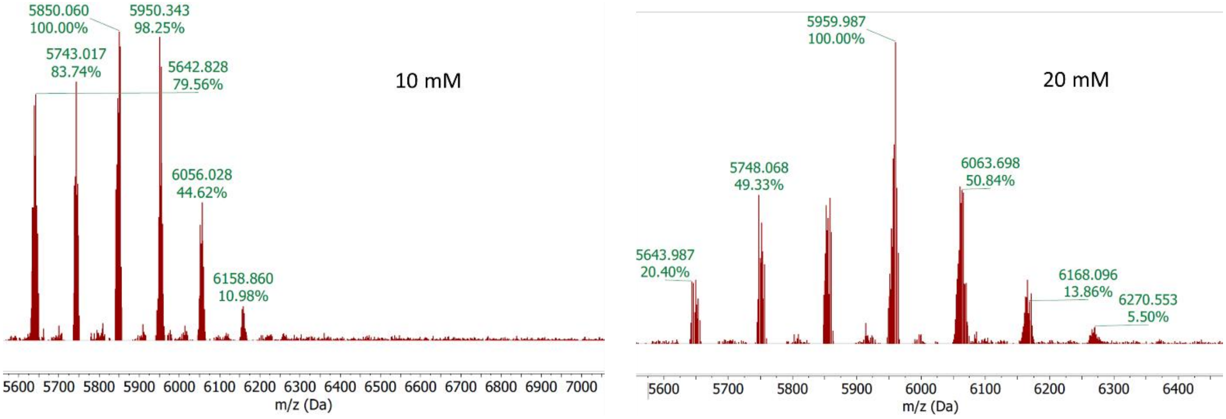


**Figure S11**. Estimation of the effective DMAP concentration in the DNA-mediated RNA arylations. MALDI-TOF spectra for 18 nt ssRNA treated with 2 mM cyanopyrimidine **5** and varied concentrations of DMAP. The concentrations of DMAP used are denoted in the corresponding spectra. Results at 2-5 mM best approximate conversions seen with the DNA-DMAP conjugate employed at 10 μM.

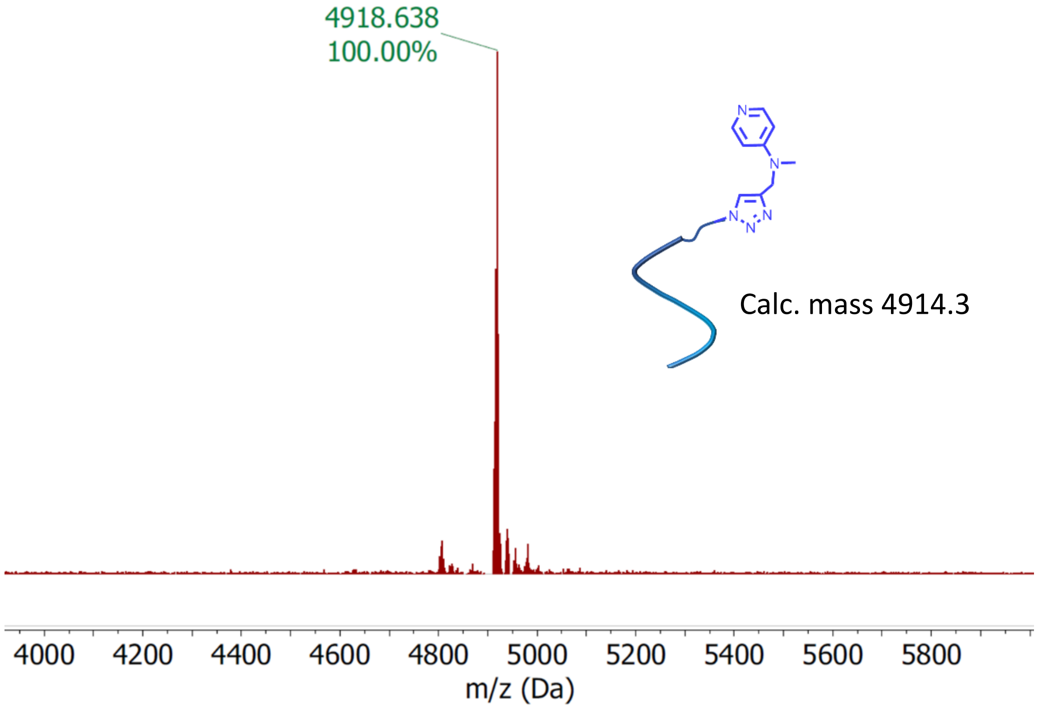


**Figure S12**. MALDI-TOF mass spectrum of DNA oligonucleotide (internal DNA-Azide**)** modified with DMAP.

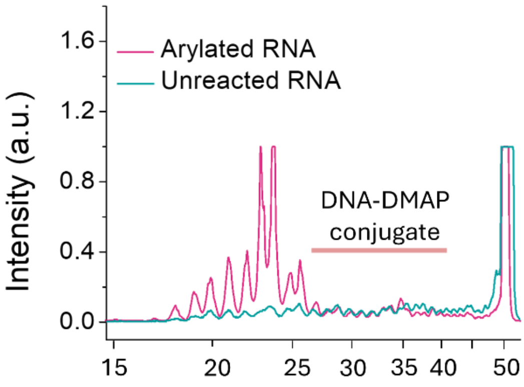


**Figure S13.** Quantification of band intensities of the lanes 3 and 4 of the gel for reaction with internal DNA-DMAP conjugate (Figure 4G); shown in x-axis are the cDNA lengths in nucleotides.

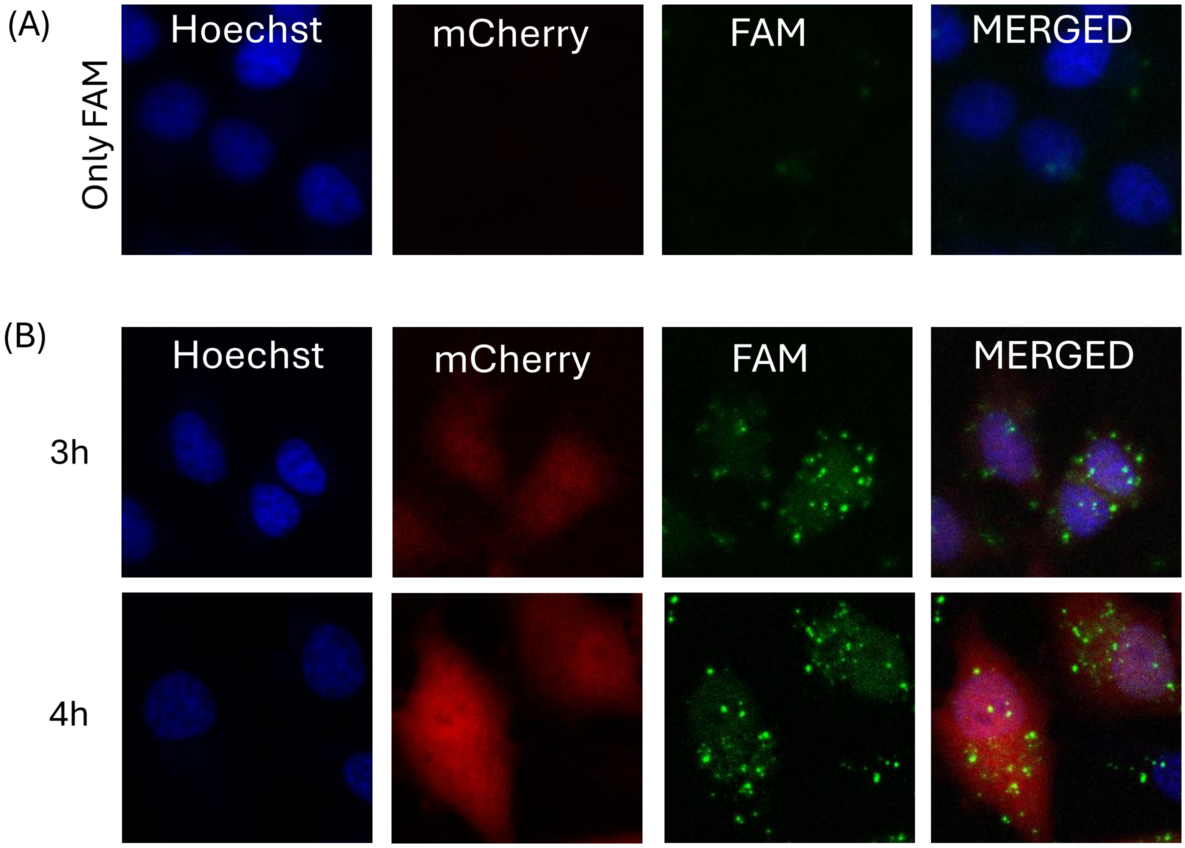


**Figure S14.** Fluorescence microscopy images of HeLa cells (A) incubated with free FAM solution (10 μM) in the absence of mCherry mRNA for 4 h, showing that green FAM signals in (B) arise from FAM conjugates of RNA; (B) early time point images of HeLa cells transfected with mCherry mRNA (3 h and 4 h) wherein the mCherry mRNA is labeled with FAM at the polyA tail prior to transfection.

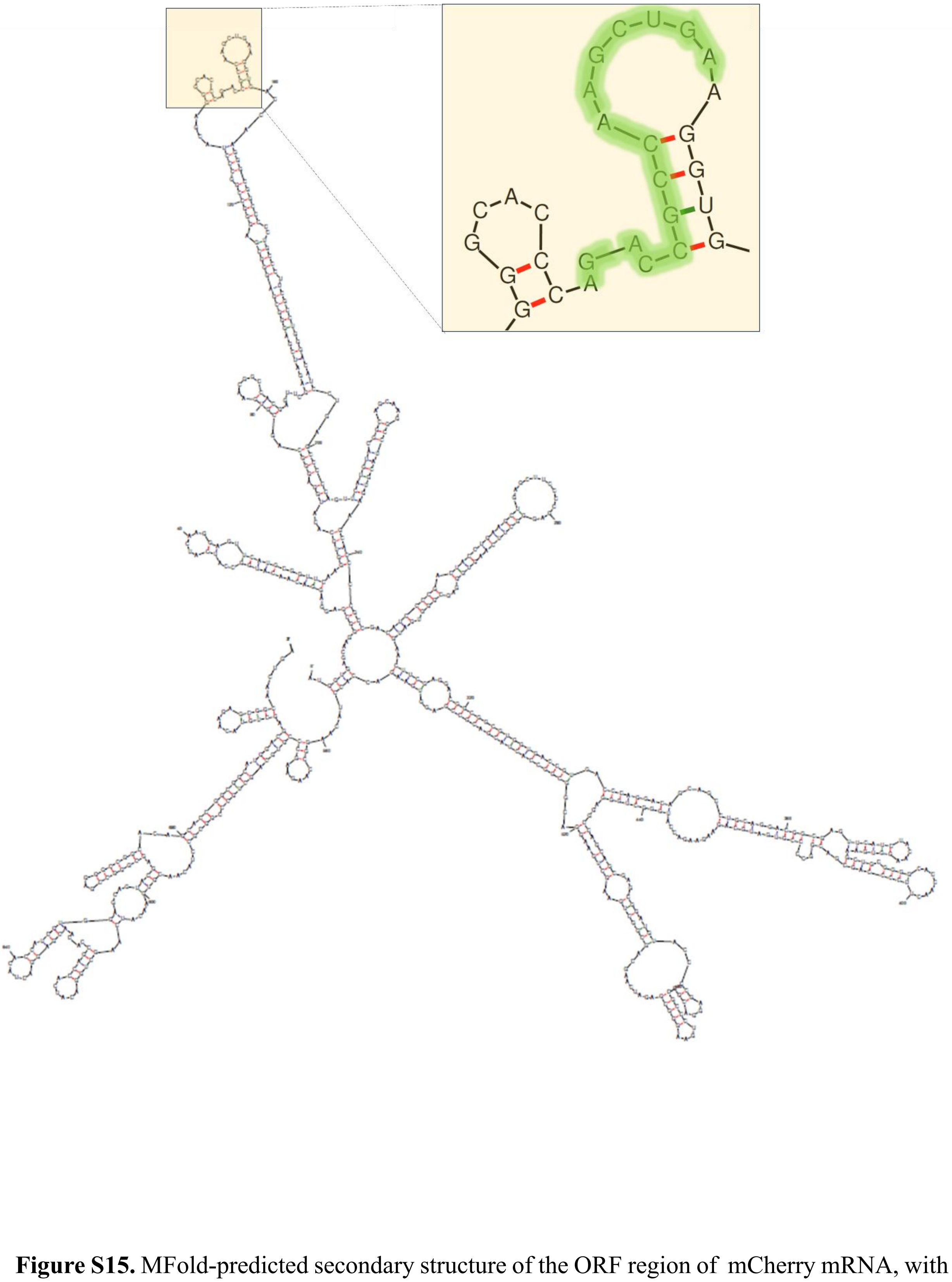

**Figure S15.** MFold-predicted secondary structure of the ORF region of mCherry mRNA, with a zoomed-in view of the targeted region. The green-highlighted segment was chosen as the target for the design of the complementary DNA-DMAP conjugate.

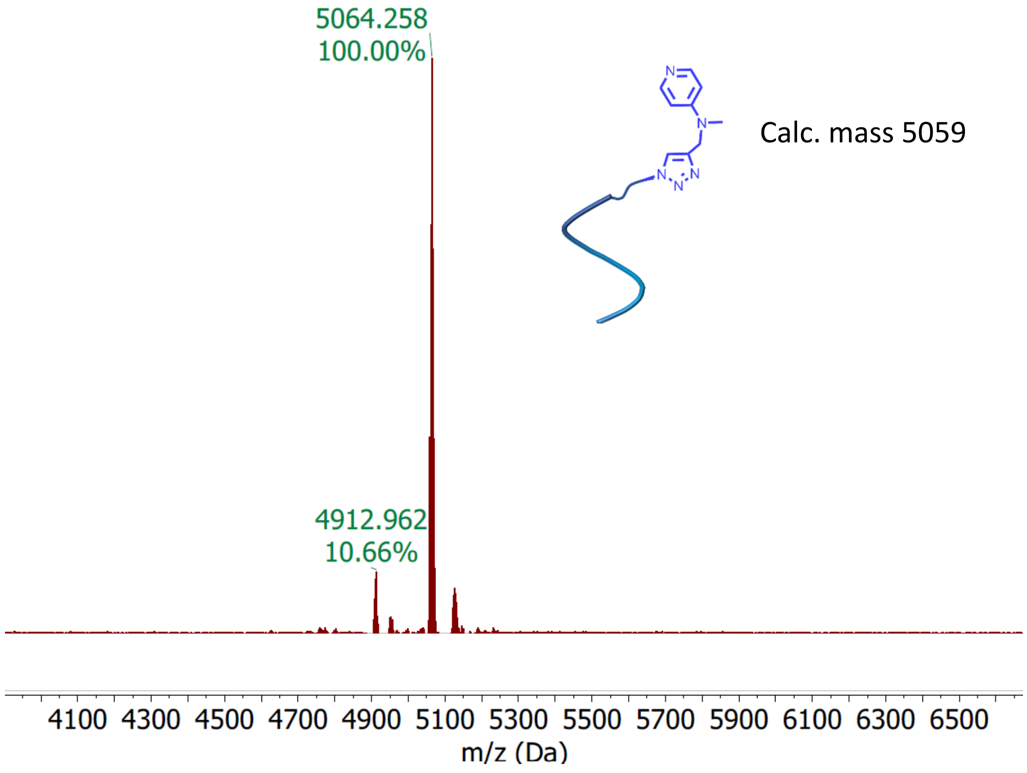


**Figure S16.** MALDI-TOF mass spectrum of DMAP-conjugated 15 nt DNA complement of mCherry mRNA.

**Uncropped PAGE gel image**

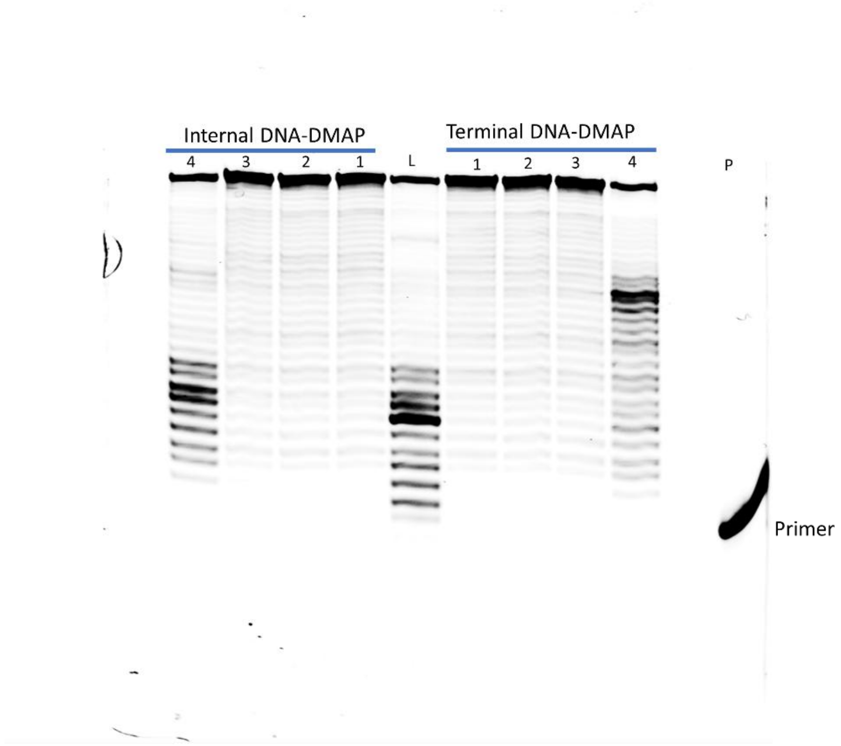


Full-size gel image for Figure 3G. “Internal DNA-DMAP” lanes in manuscript are shown as mirror image for consistency. Lane P corresponds to the primer band, and Lane L corresponds to modified 51 nt RNA, not used in the manuscript.

**Synthesis of reagents**

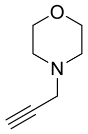


Morpholine (697 mg, 8 mmol) was weighed into a 50 mL round-bottom flask and dissolved with 10 mL DMF. Next, $K_2CO_3$ (1380 mg, 10 mmol) was added, and the reaction mixture was allowed to stir for 10 min. Then, propargyl bromide (951 mg, 8 mmol) was dissolved in 2 mL DMF and added dropwise into the reaction mixture. The reaction was allowed to stir at rt, and the progress of the reaction was monitored by TLC. After complete consumption of propargyl bromide, the reaction mixture was poured in 30 mL brine solution and extracted with ethyl acetate (3 x 30 mL). The organic layer was dried over anhydrous $MgSO_4$ and evaporated under reduced pressure. The crude mixture was purified by column chromatography using silica gel (100-200 mesh size) and hexane/ethyl acetate as the eluent (70:30, v/v). The pure product was obtained as a pale yellow liquid, Yield: 574 mg, 57%.

$^1$ H NMR (400 MHz, DMSO-*d$_6$*) δ 3.58 (t, *J* = 4.7, 2H), 3.26 (d, *J* = 2.4, 2H), 3.17 (t, *J* = 2.4, 1H), 2.43 (t, *J* = 4.7, 2H) $^{13}$C NMR (101MHz, DMSO-*d$_6$*) 79.5, 76.3, 66.5, 51.9, 46.7.

HRMS (m/z): [M+H$^+$] calcd for $C_7H_{12}NO^+$ 126.0913; found 126.0915.

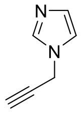

Imidazole (544 mg, 8 mmol) was weighed out into a 50 mL round-bottom flask and dissolved with 10 mL DMF. Potassium carbonate (1380 mg, 10 mmol) was added into the solution and the reaction mixture was stirred for 10 min. Next, propargyl bromide (951 mg, 8 mmol) was dissolved in 2 mL DMF and added dropwise into the reaction mixture. The reaction was stirred overnight at rt. After completion of the reaction, the reaction mixture was poured into brine solution (30 mL) and extracted with ethyl acetate (3 x 30 mL). The combined organic layer was dried over anhydrous $MgSO_4$ and evaporated under reduced pressure before purification by column chromatography using silica gel (100-200 mesh size) and hexane/ethyl acetate (65:35, v/v) as the eluent.

Imidazole propargyl: pale yellow liquid, Yield 573.7 mg, 67%

$^1$ H NMR (400 MHz, DMSO-*d$_6$*) δ 7.68 (s, 1H), 7.21 (s, 1H), 6.93 (s, 1H), 4.92 (d, *J* = 2.56, 2H), 3.51 (t, *J* = 2.56, 1H) $^{13}$C NMR (101MHz, DMSO-*d$_6$*) 137.4, 129.2, 119.7, 79.21, 76.6, 35.8; HRMS (m/z): [M+H$^+$] calcd for $C_6H_7N_2^+$ 107.0604; found 107.0409.

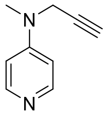

N-methyl-N-(prop-2-yn-1-yl)pyridin-4-amine,

Methyl propagylamine (276 mg, 4 mmol) was taken in a 50 mL round-bottom flask, and 8 mL DMF was added for dissolution. Nest $K_2CO_3$ (1380 mg, 10 mmol) was added, and the reaction was allowed to stir at rt for 20 min. Next, 4-Fluoropyridine (390 mg, 4 mmol), dissolved in 2 mL DMF, was added dropwise into the reaction mixture and the reaction was allowed to stir at rt for 16 h. Progress of the reaction was monitored by Thin Layer Chromatography (TLC). Following complete conversion of 4-Fluoropyridine, the reaction mixture was poured over brine solution (20 mL) and extracted thrice with ethyl acetate (3 x 30 mL). The organic layer was dried over anhydrous $Na_2SO_4$, filtered, and evaporated under reduced pressure. The crude mixture was purified by column chromatography using silica gel and hexane/ethyl acetate (70:30, v/v) as eluent. Pure product was obtained as white solid, in 86% yield (505.9 mg). $^1$ H NMR (500 MHz, DMSO-*d$_6$*) δ 8.16 (d, *J* = 5, 2 H), 6.70 (d, *J* = 5, 2 H), 4.19 (d, *J* = 2.4, 2H), 3.15 (t, *J* = 2.3, 1H), 2.95 (s, 3H). $^{13}$C NMR (125 MHz, DMSO-*d$_6$*) 153.6, 150.0, 108.4, 79.8, 75.1, 40.5, 37.4. HRMS (m/z): [M+H$^+$] calcd for $C_9H_{11}N_2^+$, 147.0917; found, 147.0919.

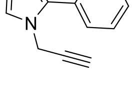

2-Phenylimidazole (720 mg, 5 mmol) was weighed into a 50 mL round-bottom flask and dissolved with 10 mL DMF. Potassium carbonate (1105 mg, 8 mmol) was added and stirred for 10 min. Propargyl bromide (594 mg, 5 mmol) was dissolved in 2 mL DMF and added dropwise into the reaction mixture. The reaction mixture was allowed to stir overnight at rt. After completion, the reaction mixture was diluted with ethyl acetate and poured over brine solution. Then the organic layer was extracted thrice with ethyl acetate (3 x 30 mL), dried over anhydrous $MgSO_4$, filtered, and evaporated under reduced pressure. The crude reaction mixture was purified by column chromatography using silica gel (100 – 200 mesh size) and hexane/ethyl acetate (70:30, v/v) as eluent.

Pale yellow liquid, Isolated yield 640.8 mg, 70 %.

$^1$H NMR (400 MHz, DMSO-$d_6$) δ 7.71 (d, $J$ = 6.8, 2H), 7.53- 7.44 (m, 3H), 7.39 (d, $J$ = 1.4, 1H), 7.04 (d, $J$ = 1.6, 1 H), 4.95 (d, $J$ = 2.5, 2 H), 3.54 (t, $J$ = 2.5, 1H). $^{13}$C NMR (101MHz, DMSO-$d_6$) 146.6, 130.8, 129.1, 129.1, 128.6, 128.6, 122.5, 79.5, 76.9, 36.8. HRMS (ESI) (m/z): $[M+H]^+$ Calcd for $C_{12}H_{11}N_2^+$: 183.0917; observed 183.0919.

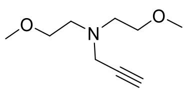

Bis (2-methoxyethyl)-amine (666 mg, 5 mmol) was weighed into a 50 mL round-bottom flask and dissolved with 10 mL DMF. $K_2CO_3$ (1105 mg, 8 mmol) was added and the solution was stirred for 10 min. Propargyl bromide (594 mg, 5 mmol) was dissolved in 2 mL DMF and added dropwise into the solution and stirred overnight at rt. After completion of reaction, the reaction mixture was diluted with ethyl acetate and poured over brine solution. The organic layer was extracted thrice (3 x 30 mL), dried over anhydrous $MgSO_4$, filtered, and evaporated under reduced pressure. The crude mixture was purified by column chromatography using silica gel (100-200 mesh size) and hexane/ethyl acetate (75:25, v/v) as eluent.

Pure compound was obtained as colourless liquid. Isolated Yield 550.8 mg, 64%.

$^1$H NMR (400 MHz, DMSO-$d_6$) δ 3.40 (m, 6 H), 3.23 (s, 6H), 3.09 (t, $J$ = 4, 1H), 2.62 (t, $J$ = 4, 2H). $^{13}$C NMR (101MHz, DMSO-$d_6$) 79.8, 75.9, 71.0, 58.4, 53.1, 43.2. HRMS (ESI) (m/z): $[M+H]^+$ Calcd for $C_9H_{17}NO_2$: 172.1332; found:172.1345.

**Sulfonyl pyrimidine with azide linker (SFPz)**

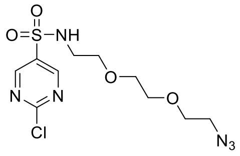

2-Chloropyrimidine-5-sulfonyl chloride (1.2 mmol, 255.6 mg) was weighed into a 50 mL round-bottom flask. 10 mL dichloromethane was added to form a suspension and stirred at 0 °C. 8-Azido-3,6-dioxaoctan-1-amine (1 mmol, 172 mg) was dissolved in 2 mL dichloromethane and added dropwise into the reaction mixture. The reaction mixture was stirred at 0 °C for 1 h, and progress of the reaction was monitored by TLC. The product was isolated by column chromatography using DCM: MeOH (98:2 v/v) as eluent. Yield 38 % (133.4 mg) as a colorless liquid.

$^{1}$ H NMR (400 MHz, DMSO-$d_6$) δ 9.11 (s, 2H), 8.27 (t, $J$ = 5.2, 1H), 3.57 (m, 2H), 3.45-3.37 (m, 8H), 3.11 (m, 2 H)

$^{13}$C NMR (101MHz, DMSO-$d_6$) 162.8, 158.8, 135.5, 70, 69.9, 69.7, 69.5, 50.4, 42.9

HRMS (ESI) (m/z): $[M+H]^+$ Calcd for $C_{10}H_{15}ClN_6O_4S$: 351.0637s; observed 351.0629.

**DMAP TC**

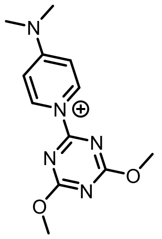


2-Chloro-4,6-dimethoxy-1,3,5-triazine (175 mg, 1 mmol) was weighed into a 20 mL glass scintillation vial equipped with a magnetic stirrer bar. 1 mL of THF was added followed by 122 mg of DMAP (1 mmol), and the reaction mixture was stirred at rt. Immediate formation of a new white precipitate was observed. After stirring for 1 h, the precipitate was filtered off and washed with ethyl acetate and diethyl ether to remove unreacted reactants. The residue was dried under vacuum to afford the pure product in quantitative yield (262 mg).

$^{1}$ H NMR (400 MHz, DMSO-$d_6$) δ 9.11, 7.25 (d, J = 8, 2H), 4.12 (s, 6H), 3.40 (s, 6H)

$^{13}$C NMR (101MHz, DMSO-$d_6$) 173.3, 164.3, 158.5, 130.8, 108.5, 56.7, 41.3

HRMS (ESI) (m/z) : $[M]^+$ Calcd for $C_{12}H_{16}N_5O_2^+$ : 262.1299, found 262.1304

**IM TC**

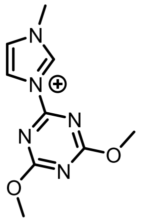


IM TC was synthetized from N-Methylimidazole and 2-Chloro-4,6-dimethoxy-1,3,5-triazine using similar synthetic protocol as DMAP TC. The product was isolated in 87 % yield (193 mg).

[1] H NMR (400 MHz, DMSO-*d6*) δ 10.38 (s, 1H), 8.47 (d, J = 1.72, 1H), 8.04 (d, J = 1.48, 1H), 4.12 (s, 6H), 4.02 (s, 3H). $^{13}$C NMR (101MHz, DMSO-*d6*) 173.4, 161.9, 138.1, 125.8, 119.4, 56.8, 37.1. HRMS (ESI) (m/z) : $[M]^+$ Calcd for $C_9H_{12}N_5O_2^+$ : 222.0986, observed 222.0989.

**DMAP CNPY**

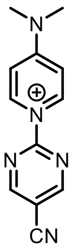


DMAP CNPY was synthesized from 2-Chloro-5-cyanopyrimidine and DMAP using similar reaction conditions as in DMAP TC. The product was obtained in quantitative yield (226 mg).

[1] H NMR (400 MHz, DMSO-*d6*) δ 9.11 (d, *J* = 8.24, 2H), 7.25 (d, *J* = 8.24, 2H), 4.12 (s, 6H), 3.40 (s, 6H). $^{13}$C NMR (101MHz, DMSO-*d6*) 163.7, 158.1, 155.5, 137.2, 115.1, 108.8, 108.2, 41.3. HRMS (ESI) (m/z) : $[M]^+$ Calcd for $C_{12}H_{12}N_5^+$ : 226.1087, observed 226.1092.

### Spectra

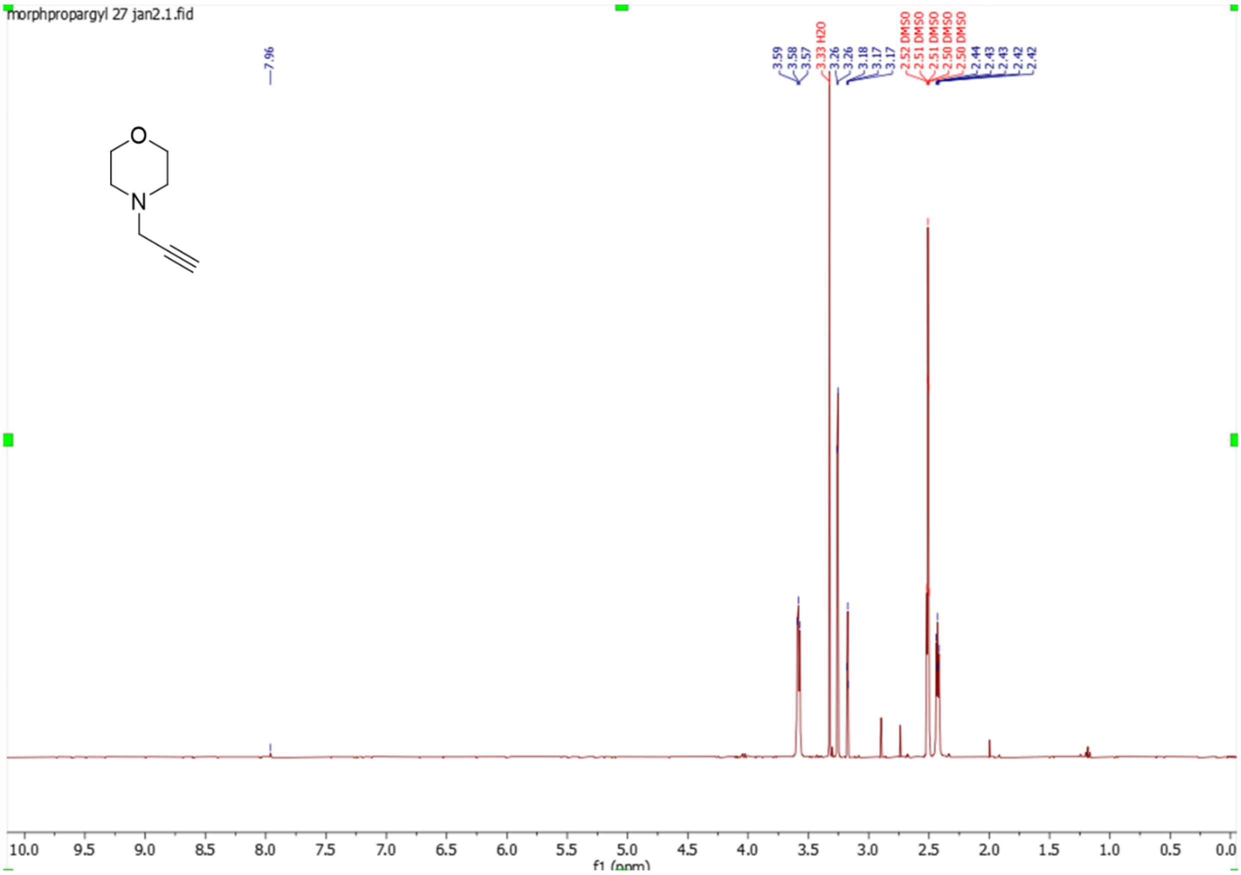
morphpropargyl 27 jan2.1.fid
7.96
3.59
3.58
3.57
3.33 H2O
3.26
3.26
3.18
3.17
3.17
2.52 DMSO
2.51 DMSO
2.51 DMSO
2.50 DMSO
2.50 DMSO
2.44
2.43
2.43
2.42
2.42
O
N
10.0
9.5
9.0
8.5
8.0
7.5
7.0
6.5
6.0
5.5
5.0
4.5
4.0
3.5
3.0
2.5
2.0
1.5
1.0
0.5
0.0
f1 (ppm)

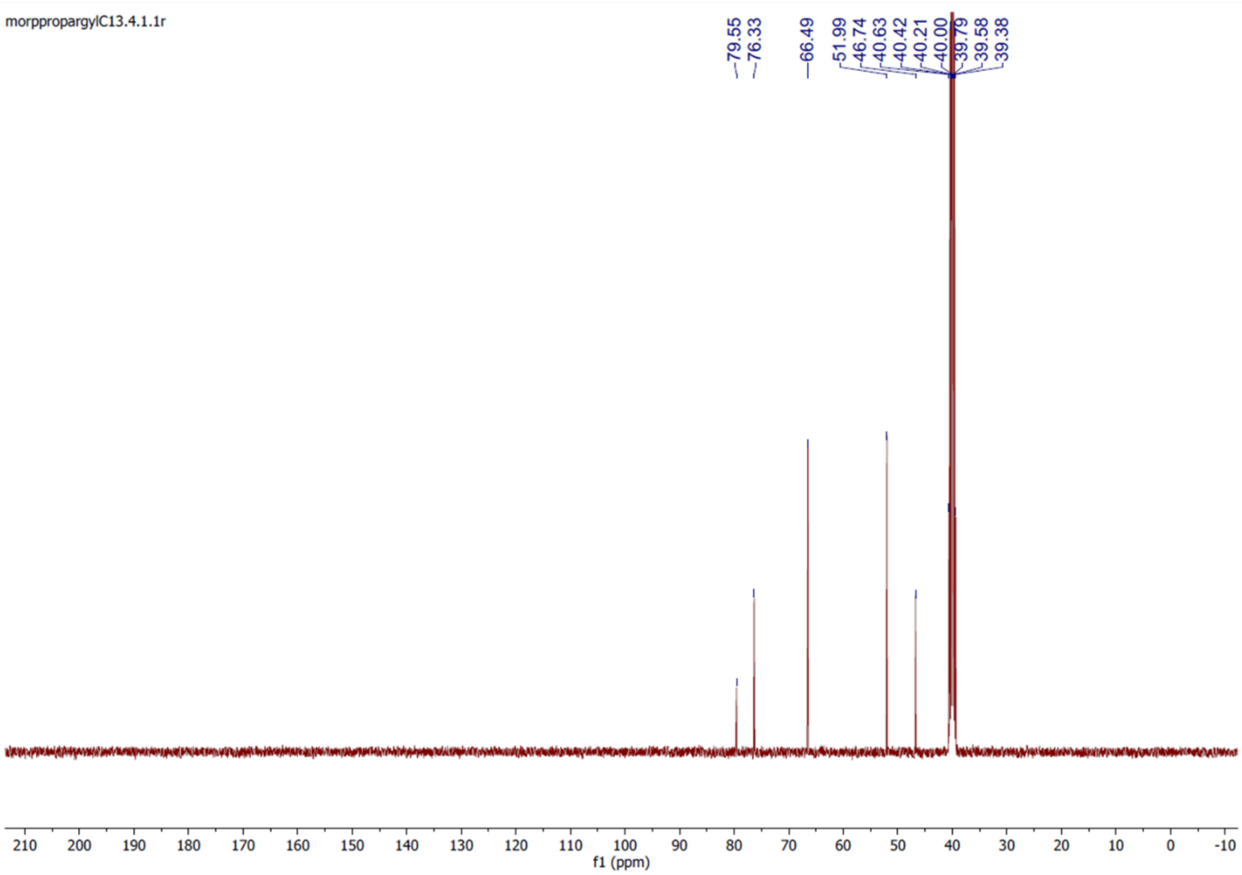
morppropargylC13.4.1.1r
79.55
76.33
66.49
51.99
46.74
40.63
40.42
40.21
40.00
39.79
39.58
39.38
210
200
190
180
170
160
150
140
130
120
110
100
90
80
70
60
50
40
30
20
10
0
-10
f1 (ppm)

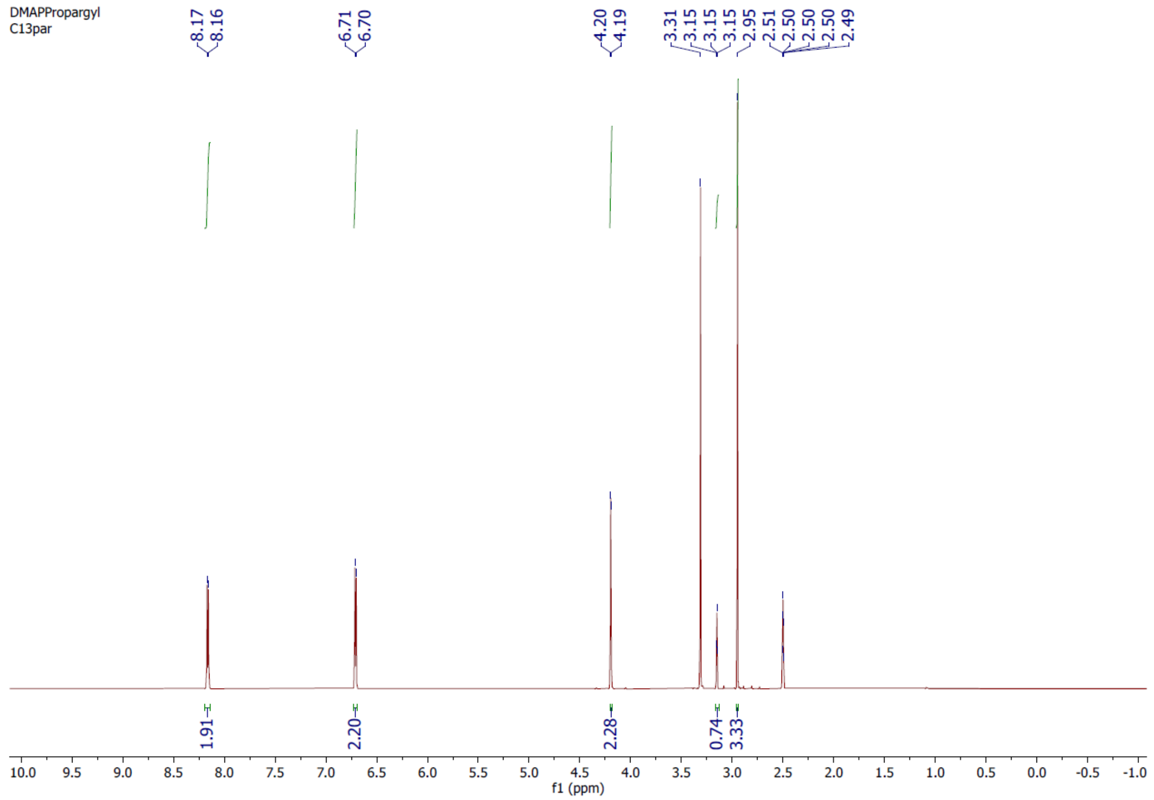
DMAPPropargyl
C13par
8.17
8.16
6.71
6.70
4.20
4.19
3.31
3.15
3.15
3.15
2.95
2.51
2.50
2.50
2.50
2.49
1.91
2.20
2.28
0.74
3.33
10.0 9.5 9.0 8.5 8.0 7.5 7.0 6.5 6.0 5.5 5.0 4.5 4.0 3.5 3.0 2.5 2.0 1.5 1.0 0.5 0.0 -0.5 -1.0
f1 (ppm)

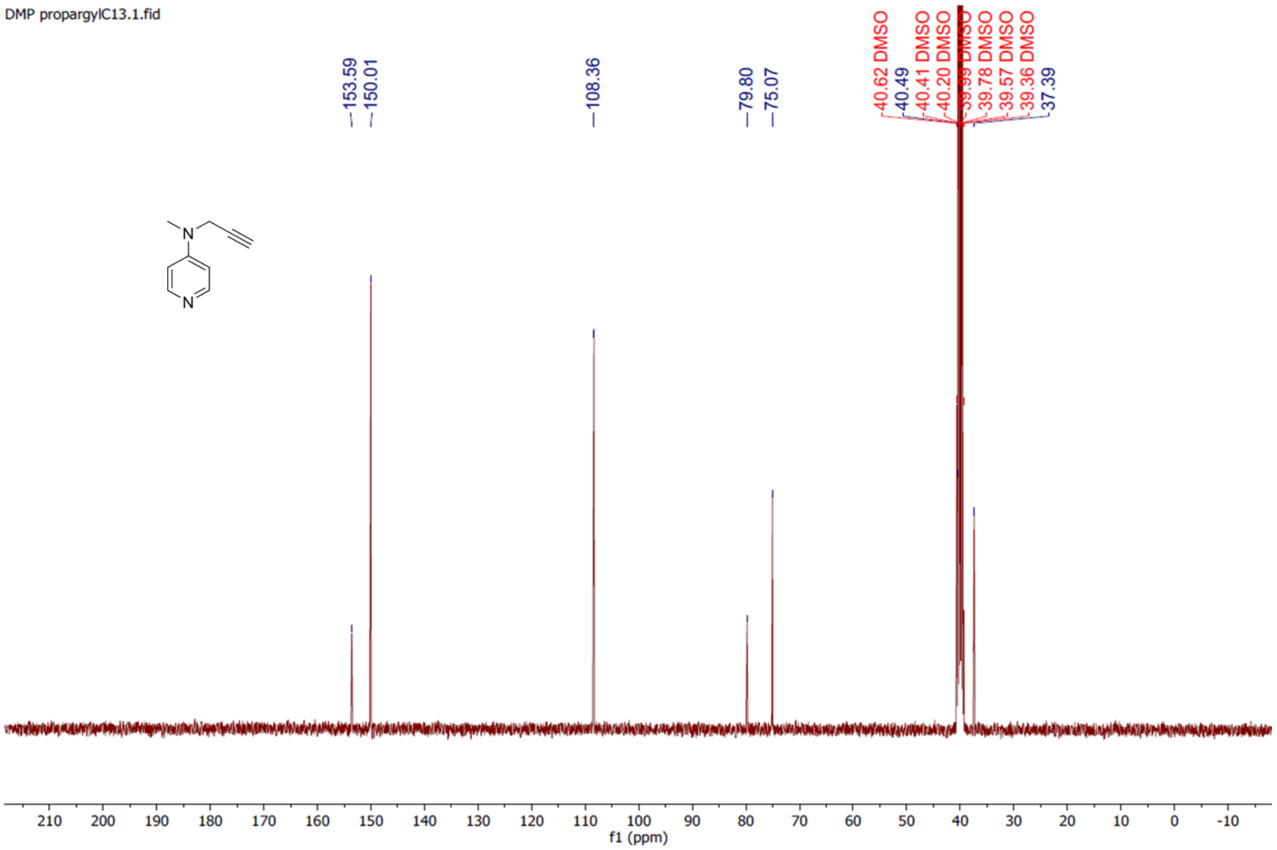
DMP propargylC13.1.fid
153.59
150.01
108.36
79.80
75.07
40.62 DMSO
40.49
40.41 DMSO
40.20 DMSO
39.99 DMSO
39.78 DMSO
39.57 DMSO
39.36 DMSO
37.39
N
N
210 200 190 180 170 160 150 140 130 120 110 100 90 80 70 60 50 40 30 20 10 0 -10
f1 (ppm)

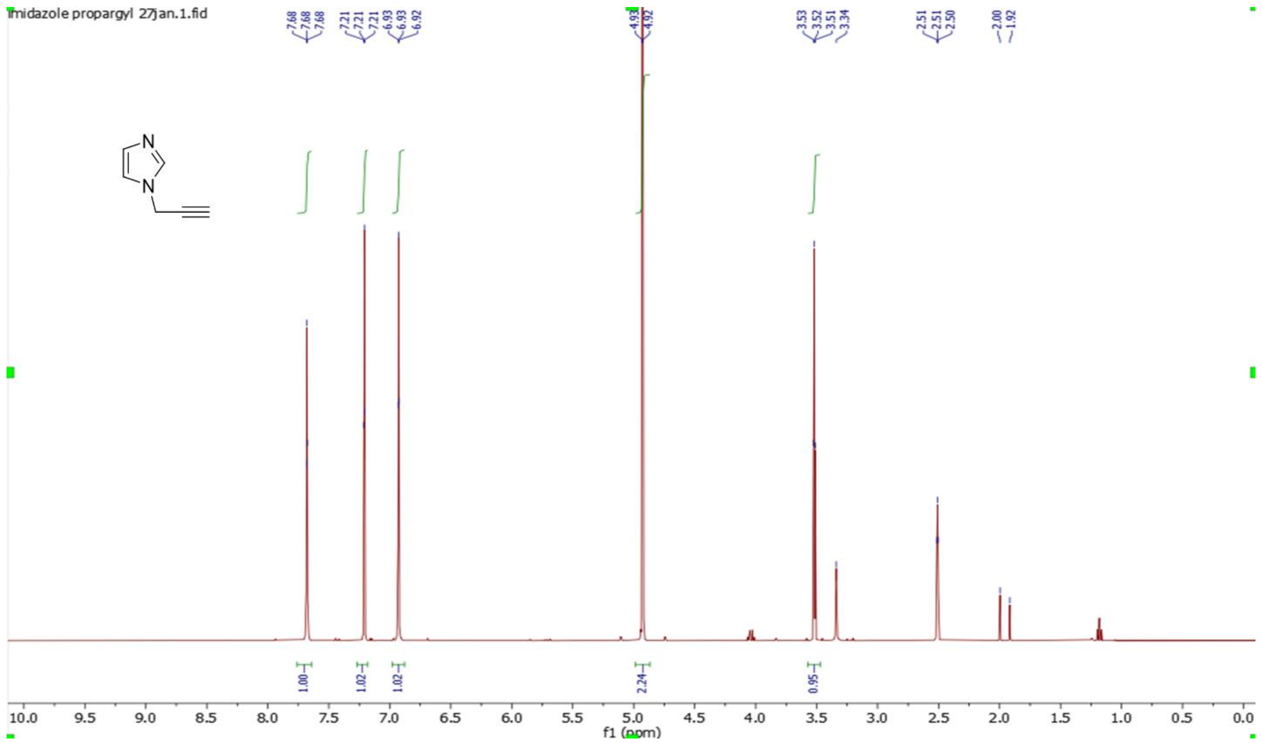
imidazole propargyl 27jan.1.fid
7.68
7.68
7.68
7.21
7.21
7.21
6.93
6.93
6.92
4.93
4.92
3.53
3.52
3.51
3.34
2.51
2.51
2.50
2.00
1.92
N
N
1.00
1.02
1.02
2.24
0.95
10.0 9.5 9.0 8.5 8.0 7.5 7.0 6.5 6.0 5.5 5.0 4.5 4.0 3.5 3.0 2.5 2.0 1.5 1.0 0.5 0.0
f1 (ppm)

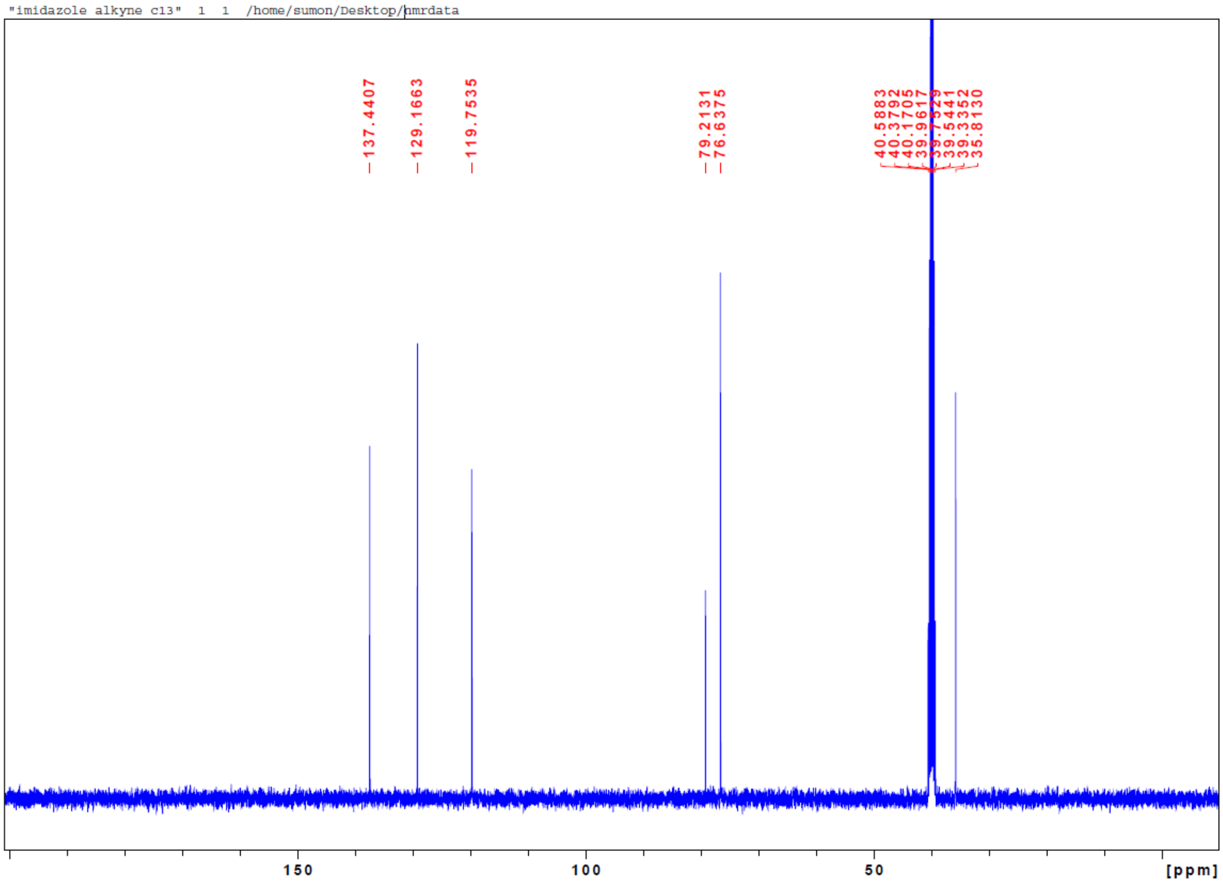
"imidazole alkyne c13" 1 1 /home/sumon/Desktop/nmrdata
137.4407
129.1663
119.7535
79.2131
76.6375
40.5883
40.3792
40.1705
39.9617
39.7529
39.5441
39.3352
35.8130
150
100
50
[ppm]

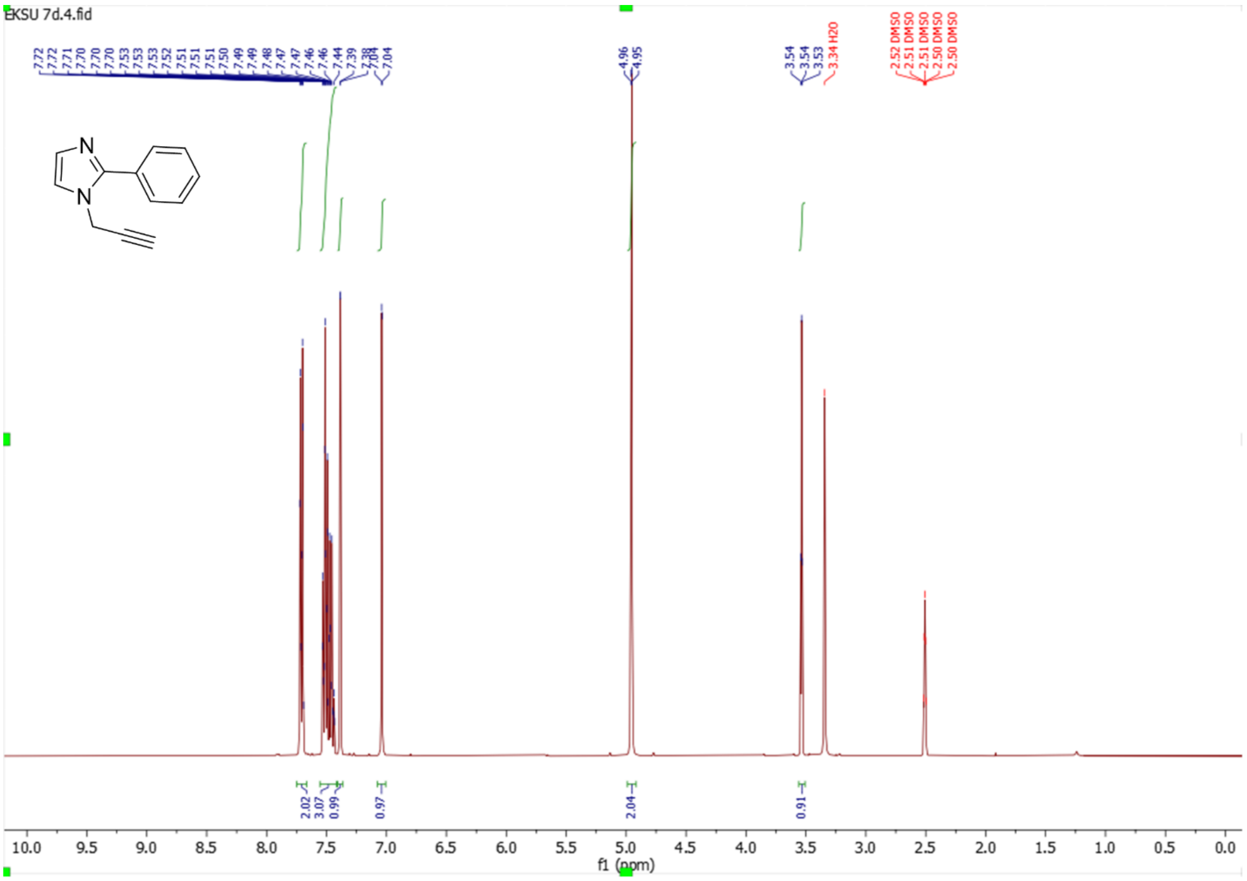
EKSU 7d.4.fid
7.72 7.72 7.71 7.70 7.70 7.70 7.53 7.53 7.53 7.52 7.51 7.51 7.51 7.50 7.49 7.49 7.48 7.47 7.47 7.46 7.46 7.44 7.39 7.38 7.04 7.04
4.96 4.95
3.54 3.54 3.53
3.34 H2O
2.52 DMSO 2.51 DMSO 2.51 DMSO 2.50 DMSO 2.50 DMSO
N
N
2.02
3.07
0.99
0.97
2.04
0.91
f1 (ppm)

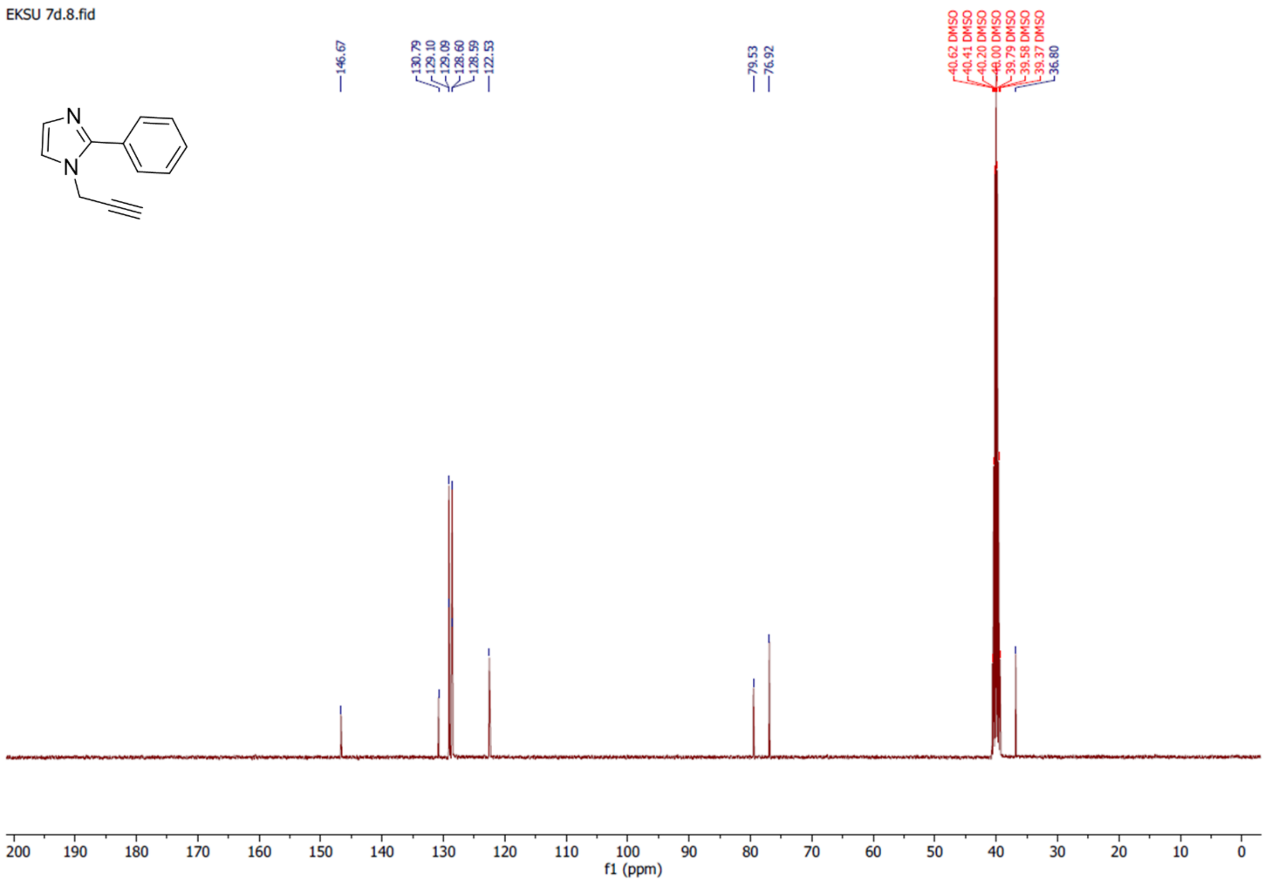
EKSU 7d.8.fid
146.67
130.79 129.10 129.09 128.60 128.59 122.53
79.53 76.92
40.62 DMSO 40.41 DMSO 40.20 DMSO 40.00 DMSO 39.79 DMSO 39.58 DMSO 39.37 DMSO 36.80
N
N
f1 (ppm)

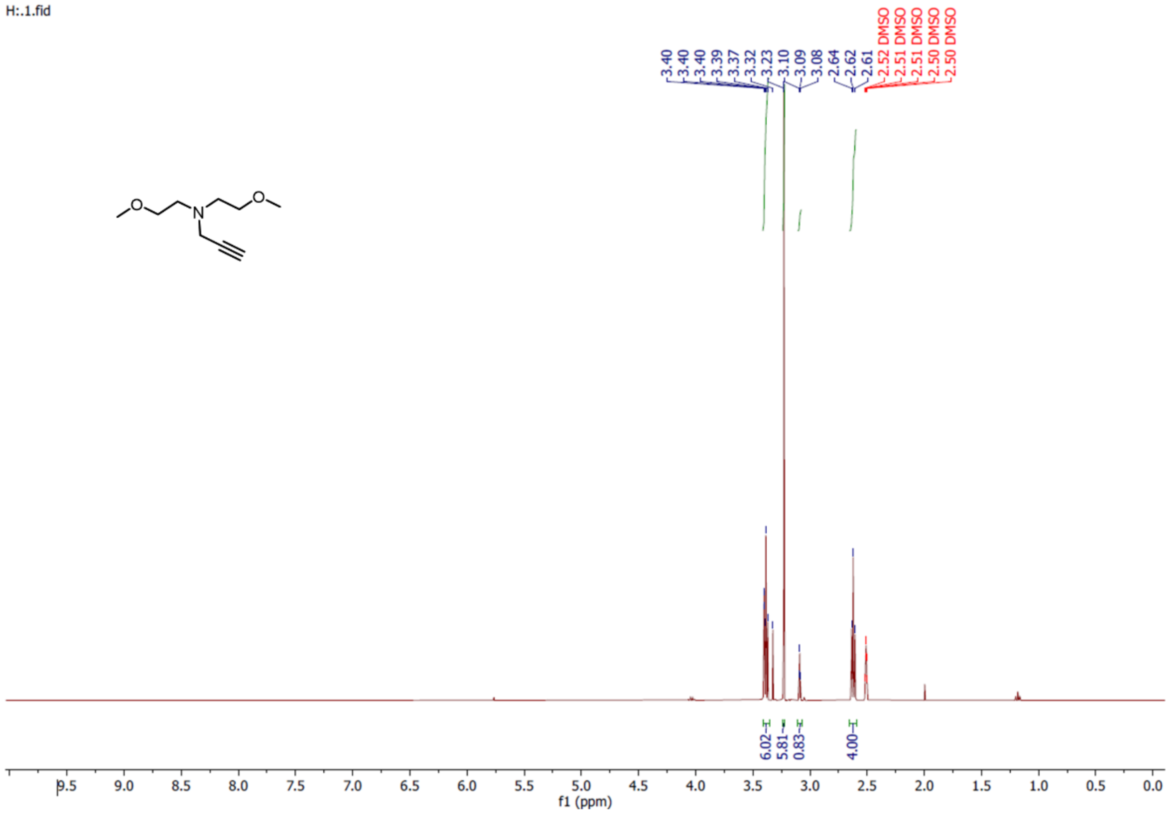
H:.1.fid
3.40
3.40
3.40
3.39
3.37
3.32
3.23
3.10
3.09
3.08
2.64
2.62
2.61
2.52 DMSO
2.51 DMSO
2.51 DMSO
2.50 DMSO
2.50 DMSO
6.02
5.81
0.83
4.00
f1 (ppm)

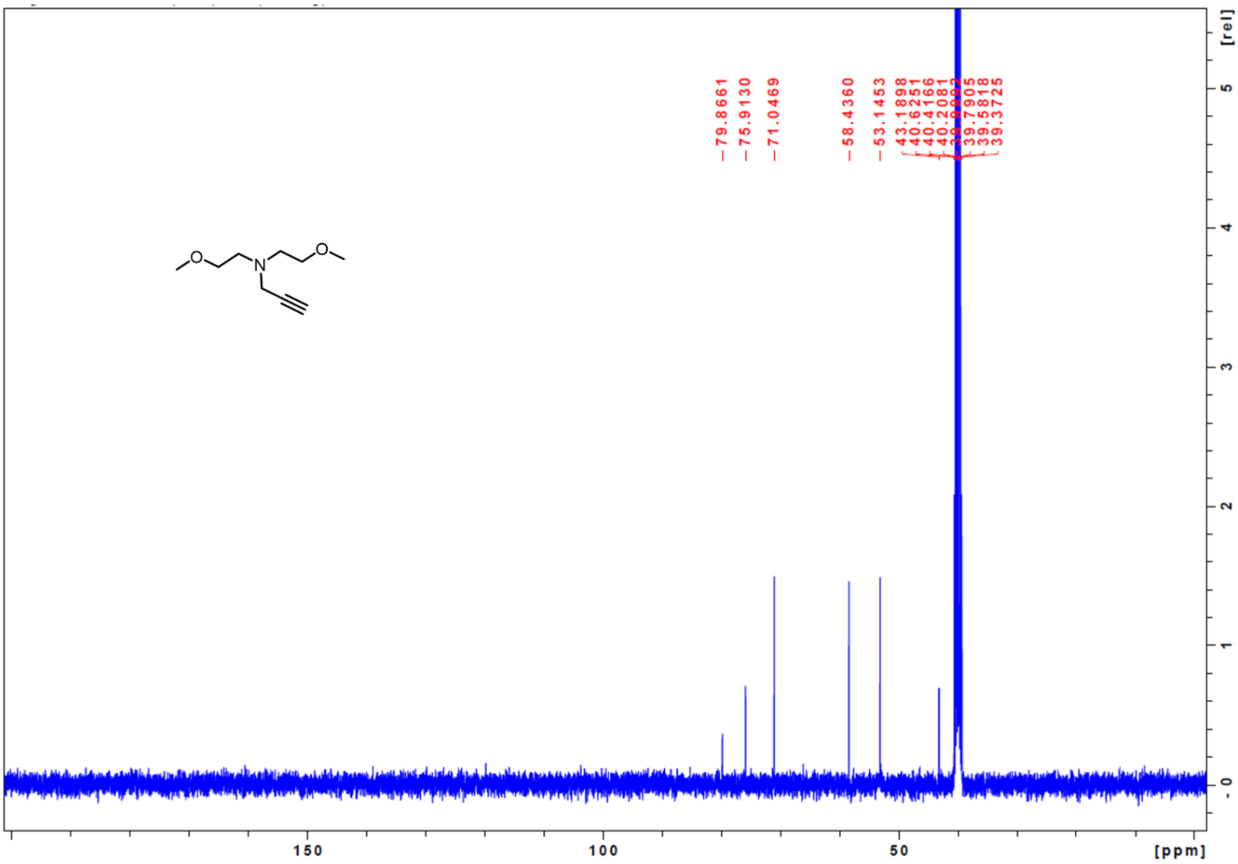
79.8661
75.9130
71.0469
58.4360
53.1453
43.1898
40.6251
40.4166
40.2081
39.7905
39.5818
39.3725
[rel]
[ppm]

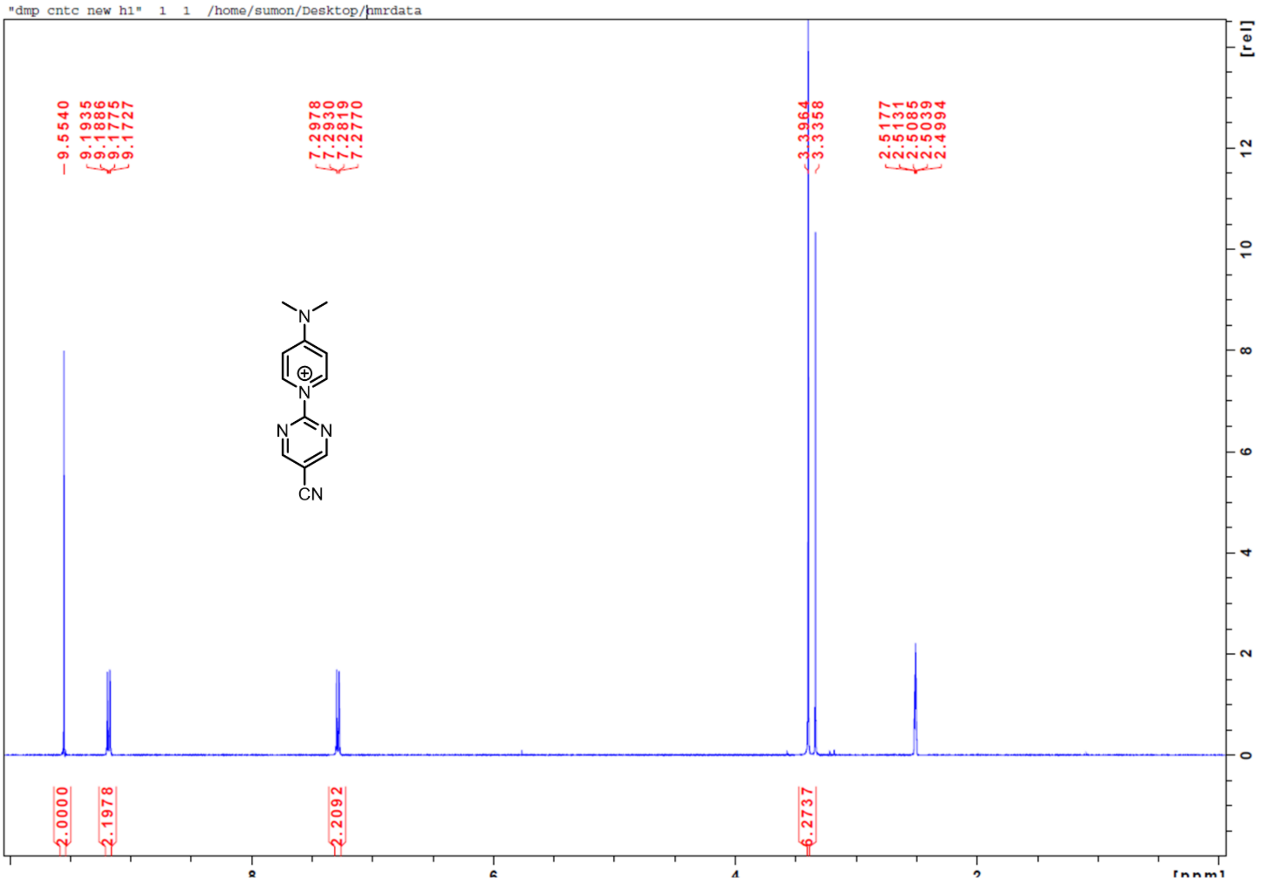
"dmp cntc new h1" 1 1 /home/sumon/Desktop/nmrdata
9.5540
9.1935
9.1886
9.1775
9.1727
7.2978
7.2930
7.2819
7.2770
3.3964
3.3358
2.5177
2.5131
2.5085
2.5039
2.4994
N
N
N
N
CN
2.0000
2.1978
2.2092
6.2737
[rel]
[ppm]

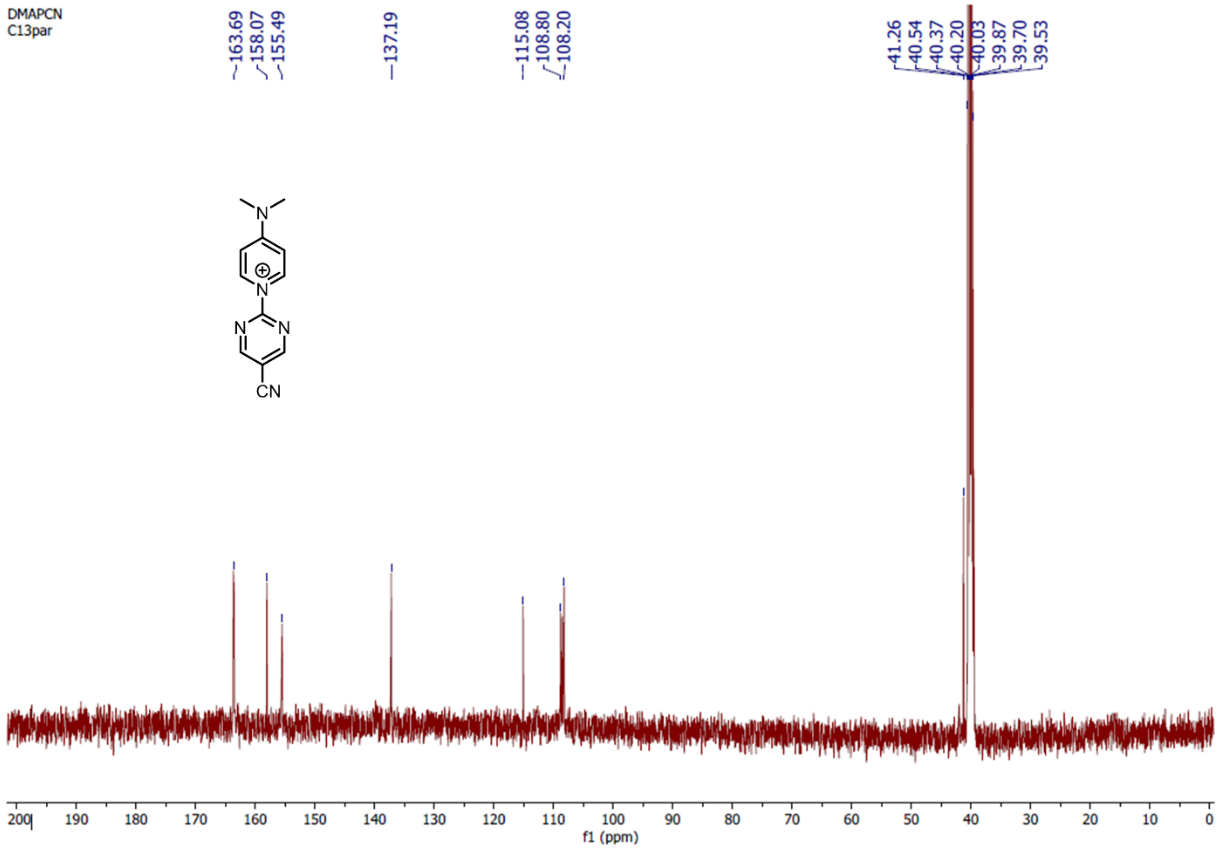
DMAPCN
C13par
163.69
158.07
155.49
137.19
115.08
108.80
108.20
41.26
40.54
40.37
40.20
40.03
39.87
39.70
39.53
N
N
N
N
CN
f1 (ppm)

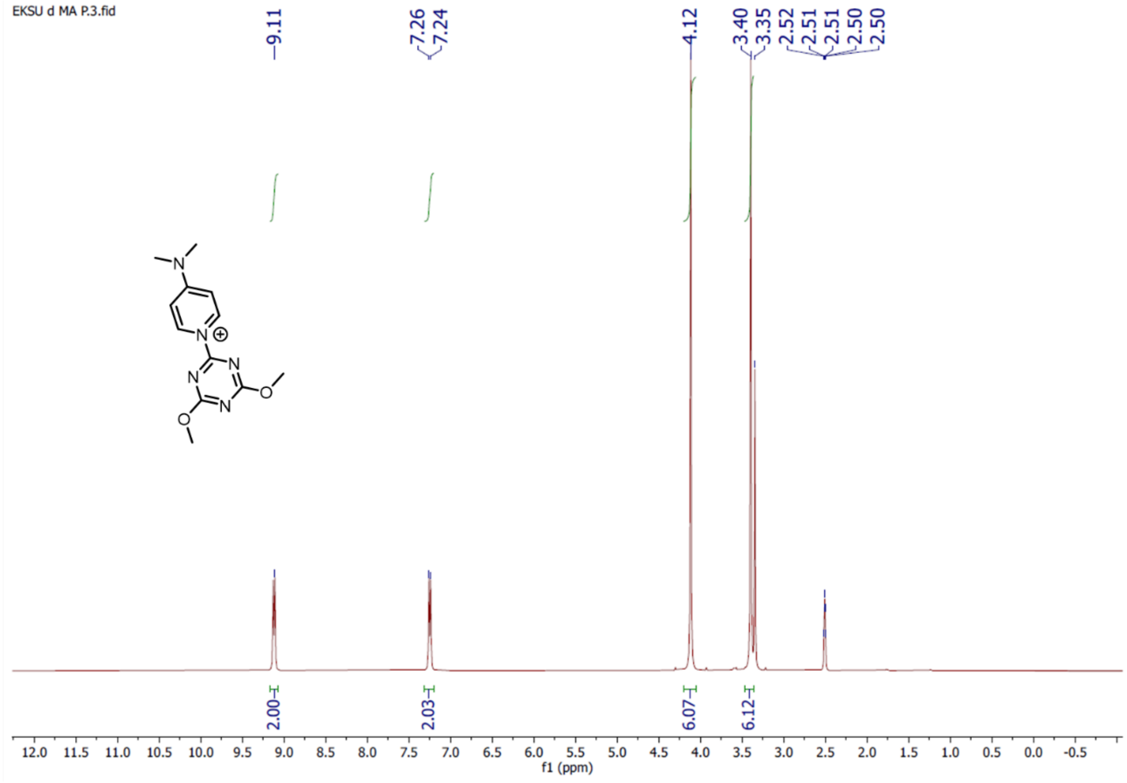
EKSU d MA P.3.fid
9.11
7.26
7.24
4.12
3.40
3.35
2.52
2.51
2.51
2.50
2.50
2.00
2.03
6.07
6.12
12.0 11.5 11.0 10.5 10.0 9.5 9.0 8.5 8.0 7.5 7.0 6.5 6.0 5.5 5.0 4.5 4.0 3.5 3.0 2.5 2.0 1.5 1.0 0.5 0.0 -0.5
f1 (ppm)

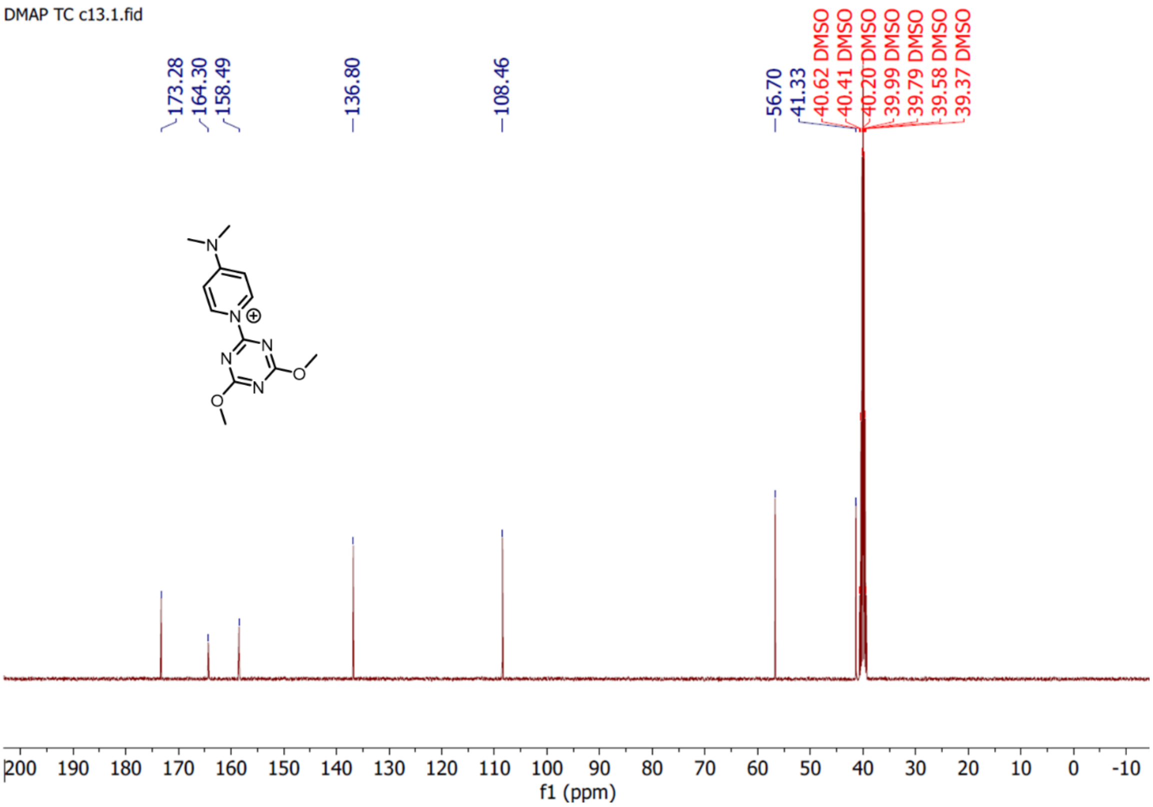
DMAP TC c13.1.fid
173.28
164.30
158.49
136.80
108.46
56.70
41.33
40.62 DMSO
40.41 DMSO
40.20 DMSO
39.99 DMSO
39.79 DMSO
39.58 DMSO
39.37 DMSO
200 190 180 170 160 150 140 130 120 110 100 90 80 70 60 50 40 30 20 10 0 -10
f1 (ppm)

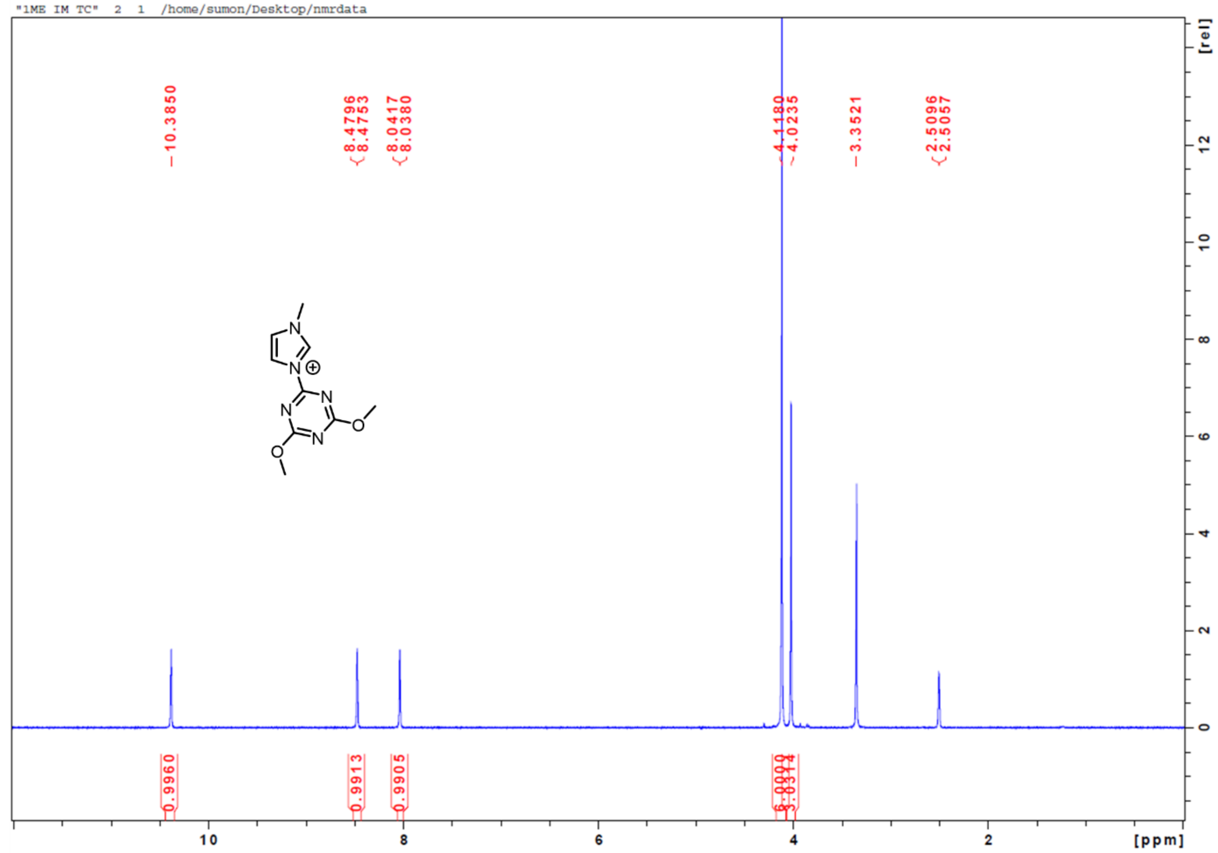
"1ME IM TC" 2 1 /home/sumon/Desktop/nmrdata
10.3850
8.4796
8.4753
8.0417
8.0380
4.1180
4.0235
3.3521
2.5096
2.5057
0.9960
0.9913
0.9905
[rel]
[ppm]

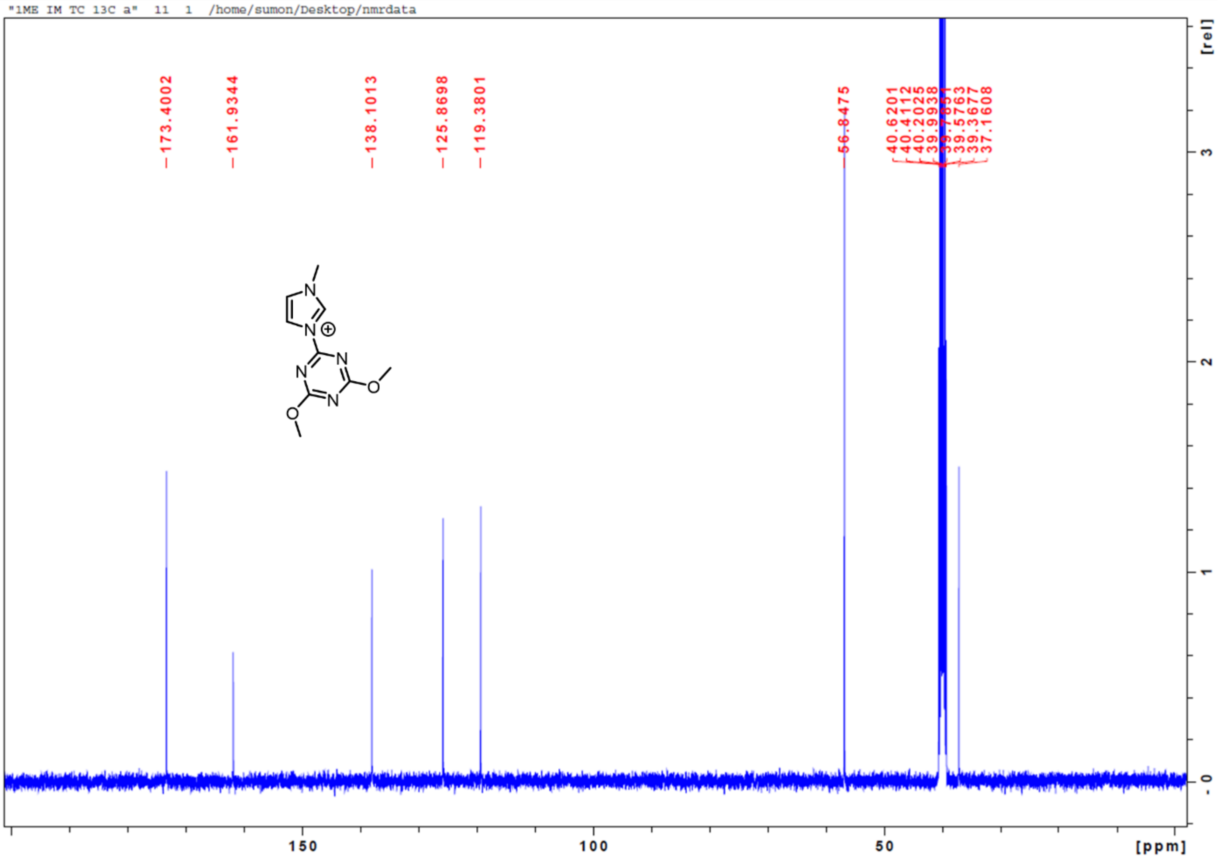
"1ME IM TC 13C a" 11 1 /home/sumon/Desktop/nmrdata
173.4002
161.9344
138.1013
125.8698
119.3801
56.8475
40.6201
40.4112
40.2025
39.9938
39.5763
39.3677
37.1608
[rel]
[ppm]

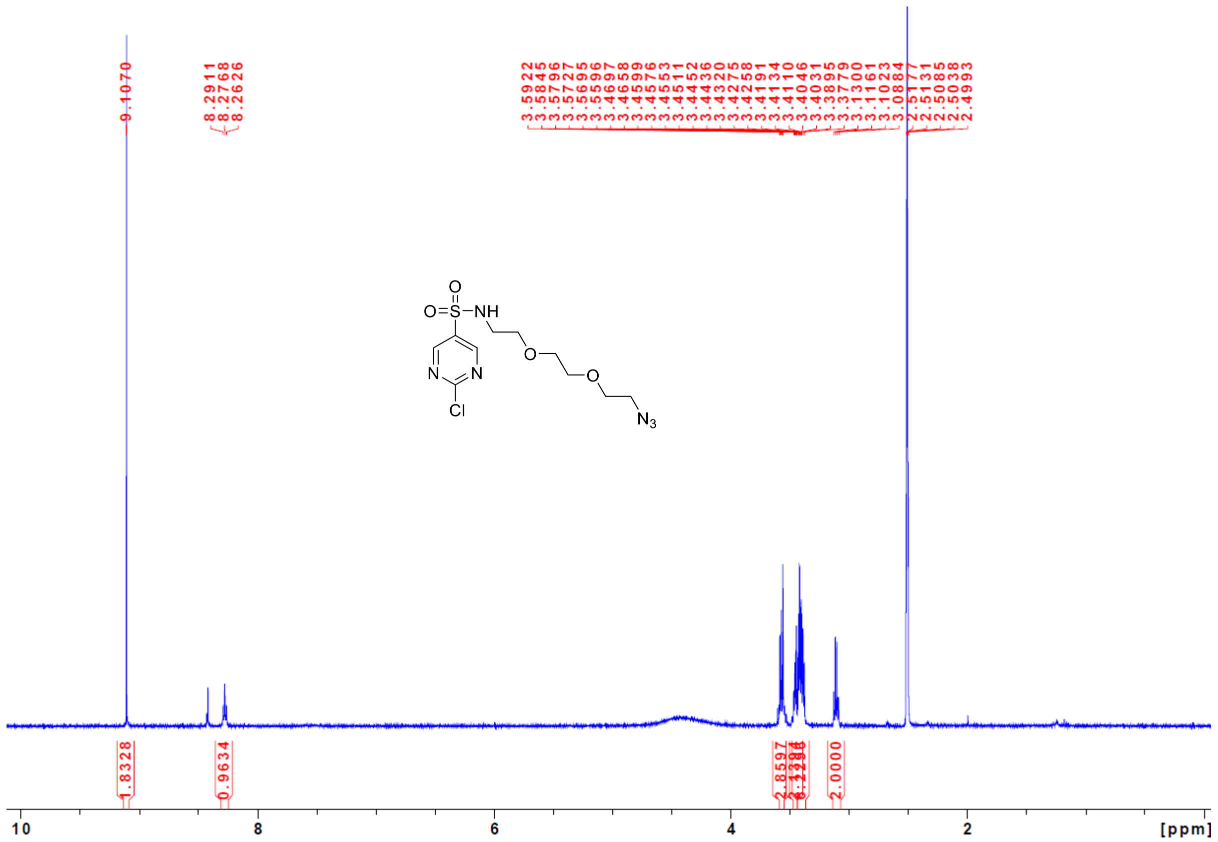

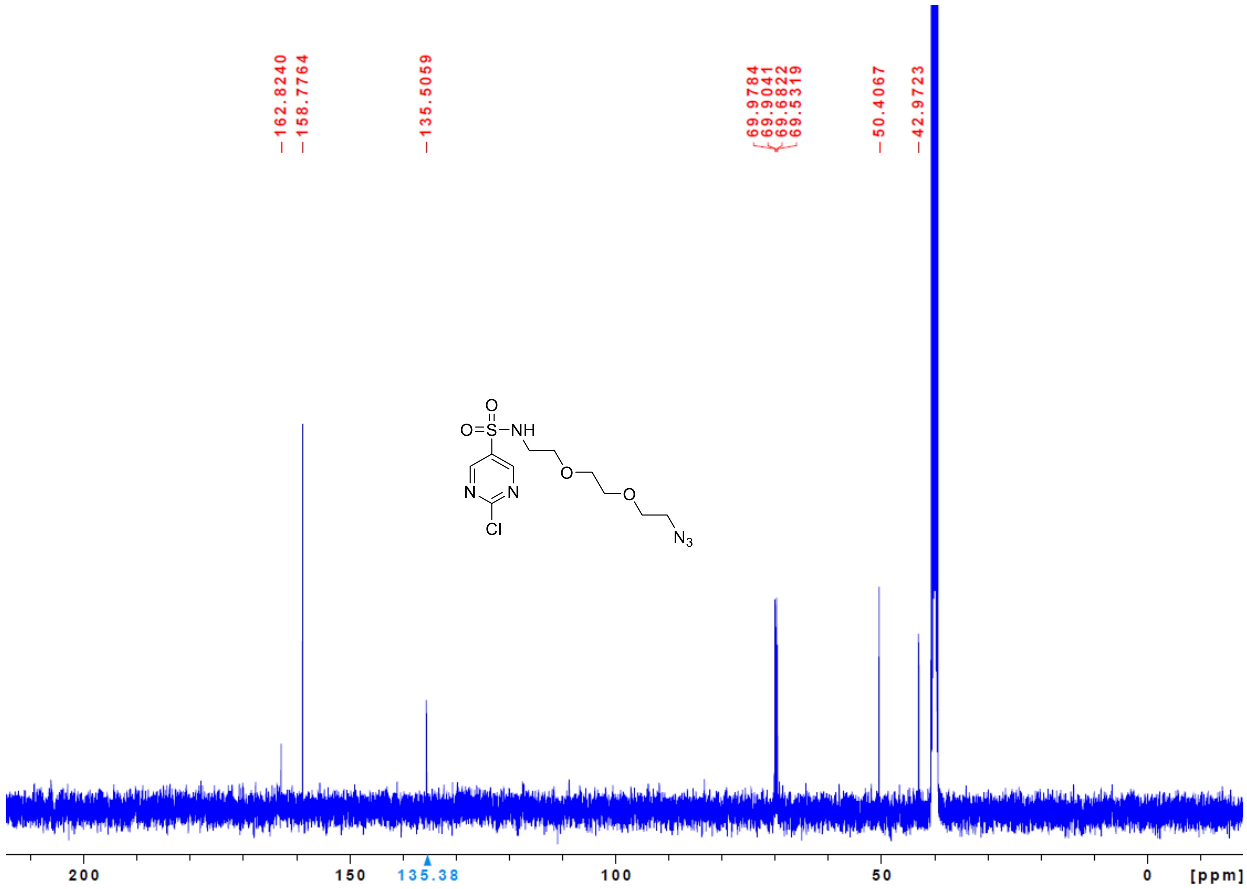
